# Altered White Matter Microstructural Organization in Post-Traumatic Stress Disorder across 3,049 Adults: Results from the PGC-ENIGMA PTSD Consortium

**DOI:** 10.1101/677153

**Authors:** Emily L Dennis, Seth G Disner, Negar Fani, Lauren E Salminen, Mark Logue, Emily K Clarke, Courtney C Haswell, Christopher L Averill, Lee A Baugh, Jessica Bomyea, Steven E Bruce, Jiook Cha, Kyle Choi, Nicholas D Davenport, Maria Densmore, Stefan du Plessis, Gina L Forster, Jessie L Frijling, Atilla Gönenc, Staci Gruber, Daniel W Grupe, Jeffrey P Guenette, Jasmeet Hayes, David Hofmann, Jonathan Ipser, Tanja Jovanovic, Sinead Kelly, Mitzy Kennis, Philipp Kinzel, Saskia BJ Koch, Inga Koerte, Sheri Koopowitz, Mayuresh Korgaonkar, John Krystal, Lauren AM Lebois, Gen Li, Vincent A Magnotta, Antje Manthey, Geoffrey J May, Deleene S Menefee, Laura Nawijn, Steven M Nelson, Richard WJ Neufeld, Jack B Nitschke, Daniel O’Doherty, Matthew Peverill, Kerry Ressler, Annerine Roos, Margaret A Sheridan, Anika Sierk, Alan Simmons, Raluca M Simons, Jeffrey S Simons, Jennifer Stevens, Benjamin Suarez-Jimenez, Danielle R Sullivan, Jean Théberge, Jana K Tran, Leigh van den Heuvel, Steven JA van der Werff, Sanne JH van Rooij, Mirjam van Zuiden, Carmen Velez, Mieke Verfaellie, Robert RJM Vermeiren, Benjamin SC Wade, Tor Wager, Henrik Walter, Sherry Winternitz, Jonathan Wolff, Gerald York, Ye Zhu, Xi Zhu, Chadi G Abdallah, Richard Bryant, Judith K Daniels, Richard J Davidson, Kelene A Fercho, Carol Franz, Elbert Geuze, Evan M Gordon, Milissa L Kaufman, William Kremen, Jim Lagopoulos, Ruth A Lanius, Michael J Lyons, Stephen R McCauley, Regina McGlinchey, Katie A McLaughlin, William Milberg, Yuval Neria, Miranda Olff, Soraya Seedat, Martha Shenton, Scott R Sponheim, Dan J Stein, Murray B Stein, Thomas Straube, David F Tate, Nic JA van der Wee, Dick J Veltman, Li Wang, Elisabeth A Wilde, Paul M Thompson, Peter Kochunov, Neda Jahanshad, Rajendra A Morey

## Abstract

A growing number of studies have examined alterations in white matter organization in people with posttraumatic stress disorder (PTSD) using diffusion MRI (dMRI), but the results have been mixed, which may be partially due to relatively small sample sizes among studies. Altered structural connectivity may be both a neurobiological vulnerability for, and a result of, PTSD. In an effort to find reliable effects, we present a multi-cohort analysis of dMRI metrics across 3,049 individuals from 28 cohorts currently participating in the PGC-ENIGMA PTSD working group (a joint partnership between the Psychiatric Genomics Consortium and the Enhancing NeuroImaging Genetics through Meta-Analysis consortium). Comparing regional white matter metrics across the full brain in 1,446 individuals with PTSD and 1,603 controls (2152 males/897 females) between ages 18-83, 92% of whom were trauma-exposed, we report associations between PTSD and disrupted white matter organization measured by lower fractional anisotropy (FA) in the tapetum region of the corpus callosum (Cohen’s *d*=−0.12, *p*=0.0021). The tapetum connects the left and right hippocampus, structures for which structure and function have been consistently implicated in PTSD. Results remained significant/similar after accounting for the effects of multiple potentially confounding variables: childhood trauma exposure, comorbid depression, history of traumatic brain injury, current alcohol abuse or dependence, and current use of psychotropic medications. Our results show that PTSD may be associated with alterations in the broader hippocampal network.

## Introduction

Posttraumatic stress disorder (PTSD) is a debilitating mental health condition with a lifetime prevalence varying globally between 1-9%^1^, with higher rates in women. Rates of PTSD are higher in populations exposed to greater levels of trauma, such as combat veterans^2^ and civilians in conflict zones^3^. In addition to trauma type, genetics, and other sociological, psychological, and biological factors, individual differences in brain structure and function may explain vulnerability to developing PTSD following exposure to trauma, may result from trauma, or may be exacerbated by PTSD^4^. Diffusion MRI (dMRI) is able to model white matter tracts and assess microstructural organization^5^. Fractional anisotropy (FA) is the most commonly used metric of microstructural organization, reflecting the degree to which water is diffusing along the axon (axially) as compared with across it (radially). Greater FA can reflect higher myelination, axonal diameter, or fiber density. Mean diffusivity (MD) reflects the average magnitude of diffusion across all directions, axial diffusivity (AD) is diffusion along the primary eigenvector (the dominant fiber direction), and radial diffusivity (RD) estimates diffusion perpendicular to the primary eigenvector. Altered microstructural organization is associated with several different psychiatric disorders and could constitute a risk factor and/or a consequence of the disorders.

There is a lack of mechanistic evidence on the effects of stress and trauma on white matter structure. Exposure to trauma could lead to white matter damage, as excessive glucocorticoid levels can be neurotoxic and can impact myelination^6,7^. Studies of white matter microstructure in PTSD have reported inconsistent results. The majority report that PTSD is associated with lower FA^8–24^, but some report higher FA^25–31^, higher and lower FA in different regions^32^, or null results^33–35^. Alterations in the cingulum bundle are frequently reported^9–13,16,18,21,23–29,31,32,36^, with differences also observed in the uncinate, corpus callosum, and *corona radiata*^14,16,18,19,24,26,29^. Inconsistent findings may be partially due to the use of hypothesis-driven rather than whole brain approaches, choice of analytic pipeline, selection of diffusion metrics, gender-specific studies, homogeneity of single cohort samples such as trauma-exposed vs. unexposed controls, and focus on military vs. civilian samples.

The PGC-ENIGMA PTSD working group is an international collaborative effort of the Psychiatric Genomics Consortium and the Enhancing NeuroImaging Genetics through Meta-Analysis (ENIGMA) consortium that aims to increase statistical power through meta- and mega-analyses of PTSD neuroimaging biomarkers. This collaborative approach has led to the largest PTSD neuroimaging study to date, reporting smaller hippocampal volume in PTSD^37^. Here, we applied this approach to investigate the microstructural organization of white matter in PTSD. The ENIGMA DTI workflow^38^, which has successfully identified white matter compromise in schizophrenia^39^, bipolar disorder^40^, major depression^41^, and 22q11.2 deletion syndrome^42^, among others, was used by 28 cohorts to process their DTI data locally. We hypothesized the largest effects of compromised microstructure will be evident in the fronto-limbic tracts, such as the cingulum, uncinate, fornix, and corpus callosum; these tracts are strongly implicated in behavioral deficits of PTSD such as emotion regulation, working memory, and episodic memory^10,16,19,20,24,43^.

## Materials and Methods

### Study Samples

The PGC-ENIGMA PTSD DTI analysis included 28 cohorts from 7 countries totaling 1,603 healthy controls and 1,446 individuals with PTSD (either formally diagnosed or with CAPS-4>40, see **Supplementary Figure 1**). The age range across cohorts was 18-83 years; all but two older Vietnam era cohorts had an average age between 29-50. Of the 3,049 participants included in these analyses, 2,073 (68%) were from military cohorts, which resulted in a disproportionate number of males (70%). The majority of cohorts included trauma-exposed controls (e.g., combat, community violence, intimate partner violence, N=1,603), although some included trauma-unexposed controls (N=122), and one included no control group. **Table 1** contains demographic and clinical information for each cohort. All participants provided written informed consent approved by local institutional review boards. Quality control was completed by each site, with visual quality checking and outlier detection. Details on ENIGMA-DTI methods^39^, inclusion/exclusion criteria, and clinical information may be found in **Supplementary Note 1, Supplementary Table 1**, and **Supplementary Note 2**, respectively.

**Table 1.** Demographic information on adult cohorts included in analyses.

| Site | Total N | M | F | N PTSD | N Control | Age range | Average age | PTSD scale | Depression scale | Type of controls | Dataset |
| --- | --- | --- | --- | --- | --- | --- | --- | --- | --- | --- | --- |
| ADNI-DoD | 134 | 134 | 0 | 70 | 64 | 61-83 | 69.3 | CAPS-4 | GDS | exposed | Military |
| Beijing | 67 | 32 | 35 | 32 | 35 | 37-61 | 49.4 | PCL-5 | na | exposed | Civilian |
| Booster | 70 | 36 | 34 | 34 | 36 | 22-59 | 39.9 | CAPS-4 | HADS-D | exposed | Police |
| Columbia | 33 | 0 | 33 | 18 | 15 | 21-53 | 33.6 | CAPS-4 | HAM-D | exposed | Civilian |
| Duke-1 | 187 | 142 | 45 | 52 | 135 | 21-57 | 39.2 | CAPS-4/5 | BDI | exposed | Military |
| Duke-2 | 88 | 61 | 27 | 36 | 52 | 23-67 | 40.0 | SCID/DTS | na | exposed | Military |
| Duke-3 | 63 | 52 | 11 | 17 | 46 | 23-65 | 38.8 | CAPS-4/5 | na | exposed | Military |
| Grady Trauma Project | 132 | 0 | 132 | 50 | 82 | 18-62 | 39.6 | CAPS-4 | BDI | exposed | Civilian |
| Groningen | 49 | 0 | 49 | 49 | 0 | 23-58 | 40.3 | CAPS-4 | BDI | no controls | Civilian |
| INTRuST | 214 | 117 | 97 | 77 | 137 | 18-56 | 36 | MINI/CAPS-4 /PCL-M/SCID | na | exposed | Military and civilian |
| iSCORE | 99 | 86 | 13 | 44 | 55 | 19-51 | 35.8 | PCL-M | CES-D | exposed | Military |
| Lawson | 98 | 52 | 46 | 46 | 52 | 18-59 | 34.7 | CAPS-4 | BDI | exposed and unexposed | Civilian |
| McLean | 55 | 0 | 55 | 41 | 14 | 18-62 | 37 | CAPS-5 | BDI | exposed | Civilian |
| Münster | 25 | 0 | 25 | 14 | 11 | 19-51 | 29 | SCID | BDI | exposed | Civilian |
| New South Wales | 162 | 62 | 100 | 85 | 77 | 18-69 | 40.2 | CAPS-4 | HAM-D | exposed | Civilian |
| South Dakota | 90 | 81 | 9 | 55 | 35 | 22-45 | 31.8 | PCL-M | na | exposed | Military |
| Stellenbosch-1 | 71 | 20 | 51 | 27 | 44 | 21-77 | 48.0 | CAPS-5 | na | both | Civilian |
| Stellenbosch-2 | 31 | 19 | 12 | 17 | 14 | 21-66 | 36.6 | CAPS-5 | na | both | Civilian |
| U Sydney | 64 | 34 | 30 | 31 | 33 | 17-49 | 36.25 | CAPS-4 | DASS | exposed | Civilian |
| UMC Utrecht | 94 | 94 | 0 | 46 | 48 | 21-57 | 35.6 | CAPS-4 | SCID | both | Military |
| UW-Madison | 48 | 44 | 4 | 25 | 23 | 22-48 | 31 | CAPS-4 | BDI | exposed | Military |
| VA Boston | 493 | 456 | 37 | 305 | 188 | 18-65 | 31.2 | CAPS-4 | na | exposed | Military |
| VA Houston | 69 | 44 | 25 | 53 | 16 | 21-58 | 31.4 | CAPS-4 | BDI | exposed | Military |
| VA Minneapolis-1 | 124 | 120 | 4 | 49 | 75 | 23-62 | 34.2 | CAPS-4 | SCID | both | Military |
| VA Minneapolis-2 | 130 | 121 | 9 | 67 | 63 | 22-59 | 32.9 | CAPS-4 | SCID | Both | Military |
| VA Waco | 53 | 46 | 7 | 36 | 17 | 25-60 | 39.6 | PCL-5 | na | unexposed | Military |
| VETSA | 239 | 239 | 0 | 33 | 206 | 56-66 | 61.8 | PCL-C | CES-D | exposed | Military |
| Yale/NCPTSD | 67 | 60 | 7 | 37 | 30 | 21-60 | 34.1 | CAPS-4 | BDI | exposed | Military |
| Overall | 3049 | 2152 | 897 | 1446 | 1603 |  | 36.9 |  |  |  |  |

### Image Acquisition and Processing

The acquisition parameters for each cohort are provided in **Supplementary Table 2**. Preprocessing, including eddy current correction, echo-planar imaging-induced distortion correction and tensor fitting, was carried out at each site. Recommended protocols and quality control procedures are available on the ENIGMA-DTI and NITRC (Neuroimaging Informatics Tools and Resources Clearinghouse) webpages. Harmonization of preprocessing schemes was not enforced across sites to accommodate site- and acquisition-specific pipelines. Once tensors were estimated, they were mapped to the ENIGMA DTI template and projected onto the ENIGMA-DTI template and were averaged within ROIs (http://enigma.ini.usc.edu/protocols/dti-protocols/). Further details and ROI abbreviations can be seen in **Supplementary Note 1**.

### Statistical Analysis

For each cohort/study, a linear model was fit using the *ppcor* and *matrixStats* packages in R 3.1.3, with the ROI FA as the response variable and PTSD and covariates as predictors. For cohorts/studies including more than one data collection site, site was included as a fixed dummy variable in the site-level analysis. As in prior ENIGMA disease working group meta-analyses^39^, a random-effects inverse-variance weighted meta-analysis was conducted at a central coordinating site (the University of Southern California Imaging Genetics Center) in R (metafor package, version 1.99– 118 http://www.metafor-project.org/) to combine individual cohort estimated effect sizes (see **Figure 2**). Cohen’s *d* for the main effect of group and unstandardized β coefficients (regression parameters) for continuous predictors were computed with 95% confidence intervals. We used the Cohen’s d calculation that accounts for covariates in the fixed effects model, using the following equation:

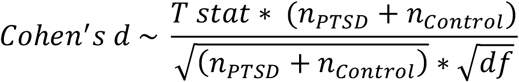

Heterogeneity scores (*I*^2^) for each test were computed, indicating the percent of total variance in effect size explained by heterogeneity across cohorts. Bilaterally averaged FA was the primary imaging measure, with corresponding MD, RD, and AD examined *post hoc* when FA was significant for an effect of diagnosis. Lateralized ROIs were examined *post hoc* when a significant association was found with the bilateral average. The corticospinal tract was not analyzed as it has poor reliability^38^. A conservative Bonferroni correction was used for multiple testing (*p*<0.05/24=0.0021; 18 bilateral ROIs, 5 midline ROIs, average FA). *Non-linear age term*: We first conducted analyses to examine whether a non-linear age term should be included in statistical models along with age and sex, as age has a non-linear effect on FA^44^. As this analysis did reveal a significant effect of non-linear age above and beyond linear age, age^2^ was included in all subsequent analyses. *Primary – group comparison*: We compared PTSD cases to all controls (both trauma-exposed and unexposed), PTSD cases to trauma-exposed controls only, and trauma-exposed to trauma un-exposed controls. *Secondary – subgroups*: We examined PTSD associations in males and females separately, and in military and civilian samples separately. These results may be found in **Supplementary Note 3**. *Secondary – interactions*: We examined potential interactions between PTSD and age or sex. These results can be seen in **Supplementary Note 4**. *Secondary – additional covariates*: We tested a model including ancestry, but as this was a meta-analysis and most cohorts were primarily composed of participants of white non-Hispanic descent, this had a very minimal impact, and we did not include this variable as a covariate in our analysis. We examined the impact of five potentially confounding covariates on the associations of PTSD with FA – childhood trauma, depression, alcohol dependence/abuse, traumatic brain injury (TBI, of any severity), and use of psychotropic medications. We compared the white matter microstructure of individuals with PTSD to that of controls with each covariate included individually in the model, and in the subset of sites that collected data on childhood trauma, depression, alcohol use disorders, TBI, or medication without that covariate in the model to determine whether differences in results were due to the inclusion of the covariate or the reduction in sample size. There were not enough participants with all five variables to simultaneously model these potential confounds in a single model. Details of these methods and results are provided in **Supplementary Note 5**. Briefly, binary variables were created for depression, TBI, and medication use. As depression was assessed using a variety of measures, we used published cut-offs to recode the data as categorical depression (see **Supplementary Note 2** for more details). Alcohol use disorders and childhood trauma were coded as three-level ordinal variables based on evidence of dose-dependent effects on brain structure and clinical severity, respectively^45,46^: Alcohol use disorders: 0=no alcohol use disorder, 1=alcohol abuse, 2=alcohol dependence, as measured by the SCID or AUDIT^47^; childhood trauma (as measured by the Childhood Trauma Questionnaire): 0=no reported childhood trauma, 1=one type of childhood trauma exposure, 2=two or more types of childhood trauma exposure; *PTSD severity*: To examine PTSD severity, we conducted linear regressions on CAPS-4 score in the PTSD group for sites that collected CAPS-4. We examined linear associations with CAPS-4 score covarying for childhood trauma, depression, alcohol use disorders, TBI, and medication use, and we tested associations with CAPS-4 separately in military veterans and civilians as well as males and females (see **Supplementary Note 5** for more details).

## Results

### Group differences

We found significantly lower FA in the PTSD group in the tapetum of the corpus callosum (*d*= −0.12, *p*=0.0021) when comparing PTSD (n=1,397) and all controls (n=1,603). *Post hoc* analysis revealed a larger effect in the left than in the right tapetum (left *d*=−0.14, *p*=0.00040; right *d*=−0.080, *p*=0.038). *Post hoc* analysis also revealed higher RD in the tapetum in the PTSD group (bilateral *d*=0.10, *p*=0.0085; left *d*=0.12, *p*=0.0016).

In the analysis comparing participants with PTSD (n=1,339) to trauma-exposed controls (n=1,481), we found lower FA and higher RD in the tapetum in PTSD, although the bilateral tapetum was marginally significant (FA: bilateral *d*=−0.11, *p*=0.0044; left *d*=−0.14, *p*=0.00065; RD: bilateral *d*=0.090, *p*=0.027; left *d*=0.12, *p*=0.0026) (see **Table 2** and **Figure 1**, and see **Figure 2** for site-specific effects). PTSD participants from cohorts that only included trauma-unexposed controls were not included.

**Figure 1.**
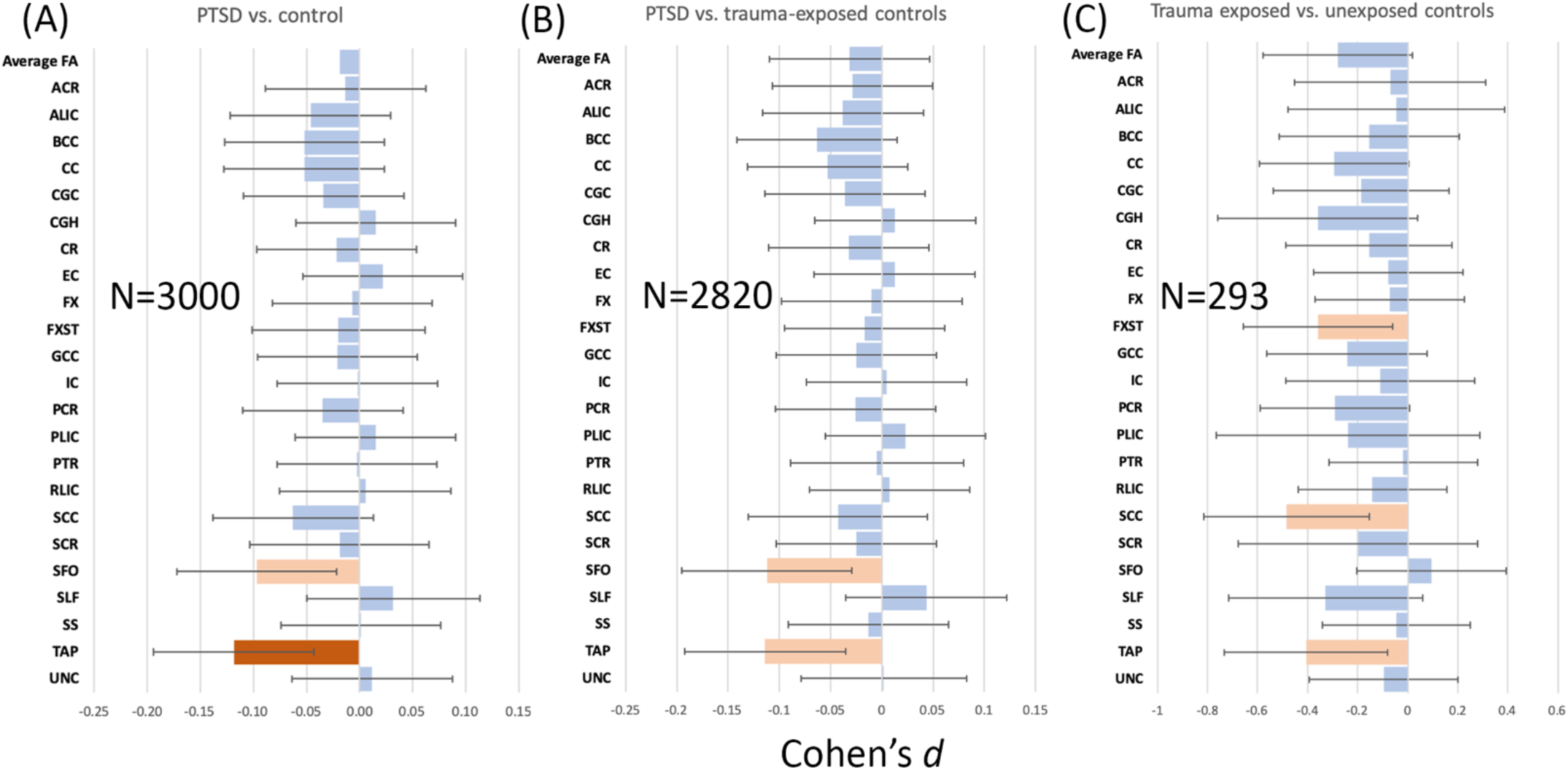
Results from the group comparisons. (A) Results comparing PTSD to all controls; (B) results comparing PTSD to trauma-exposed controls only; (C) results comparing trauma-exposed to unexposed participants. Cohen’s *d* statistics are shown across all bilateral and midline ROIs and average FA, with bars indicating the 95% confidence interval. The ROI abbreviations are explained in **Supplementary Note 1**. As PTSD was coded “1” and control “0”, negative statistics indicate lower FA in PTSD. Total N is listed for each comparison. Dark orange bars indicate significance (*p*<0.0021) and light orange bars indicates marginally significant results (0.05>*p*>0.0021).

**Figure 2.**
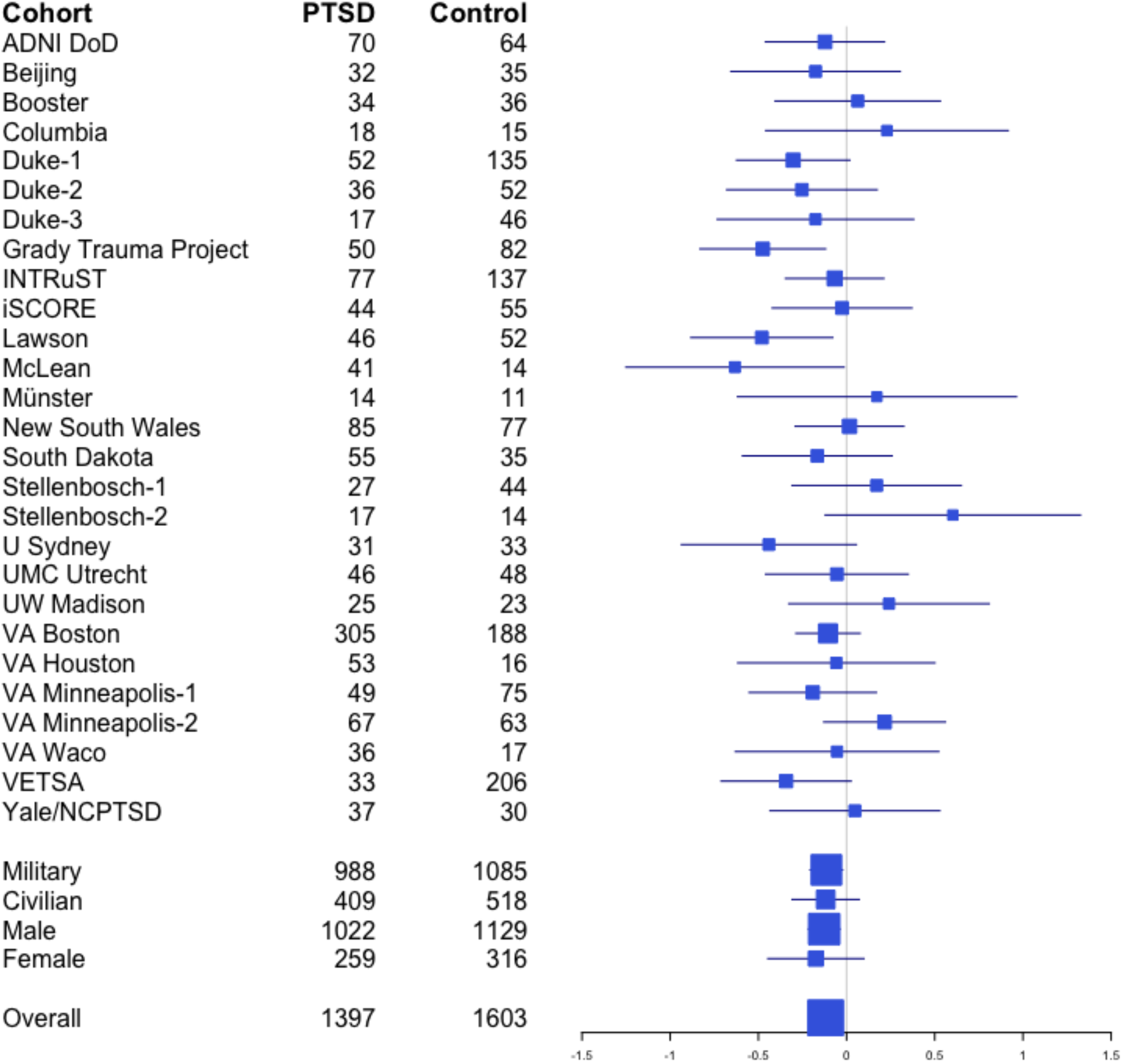
Site effects for tapetum result. Forest plot shows the effect sizes (Cohen’s *d*) for each of the 25 cohorts, scaled by sample size, with bars for 95% CI. The effect size and 95% CI of the meta-analysis is shown at the bottom of the figure, along with effect sizes and 95% CI for the subgroup analyses of the military cohorts, civilian cohorts, male cohorts, and female cohorts.

**Table 2.** Results from the group comparisons. (A) Results comparing PTSD to all controls, (B) results comparing PTSD to trauma-exposed controls only. Cohen’s *d* values, uncorrected and Bonferroni-corrected *p*-values, the 95% confidence interval for the *d* statistic, and the I^2^ (heterogeneity) are shown for the group comparisons. **Bolded** results are significant when corrected for multiple comparisons, *italicized* results are marginally significant.

| ROI | (A) PTSD vs all controls |  |  |  | (B) PTSD vs trauma-exposed controls |  |  |  |
| --- | --- | --- | --- | --- | --- | --- | --- | --- |
|  | Meta *d* | Meta *p*-value<br>uncorr./corr. | 95% CI | *I*<sup>2</sup> | Meta *d* | Meta *p*-value<br>uncorr./corr. | 95% CI | *I*<sup>2</sup> |
| Average FA | -0.02 | 0.63/0.97 | [-0.09, 0.06] | 0 | -0.03 | 0.43/0.90 | [-0.11, 0.05] | 0 |
| ACR | -0.01 | 0.73/0.97 | [-0.09, 0.06] | 0.19 | -0.03 | 0.47/0.90 | [-0.11, 0.05] | 0 |
| ALIC | -0.05 | 0.23/0.92 | [-0.12, 0.03] | 0 | -0.04 | 0.34/0.90 | [-0.12, 0.04] | 0 |
| BCC | -0.05 | 0.18/0.86 | [-0.13, 0.02] | 0 | -0.06 | 0.11/0.90 | [-0.14, 0.02] | 0 |
| CC | -0.05 | 0.18/0.86 | [-0.13, 0.02] | 0 | -0.05 | 0.18/0.90 | [-0.13, 0.03] | 0 |
| CGC | -0.03 | 0.38/0.97 | [-0.11, 0.04] | 0 | -0.04 | 0.36/0.90 | [-0.11, 0.04] | 0 |
| CGH | 0.02 | 0.69/0.97 | [-0.06, 0.09] | 0 | 0.01 | 0.74/0.95 | [-0.07, 0.09] | 0 |
| CR | -0.02 | 0.57/0.97 | [-0.10, 0.05] | 0 | -0.03 | 0.41/0.90 | [-0.11, 0.05] | 0 |
| EC | 0.02 | 0.57/0.97 | [-0.05, 0.10] | 0.01 | 0.01 | 0.75/0.95 | [-0.07, 0.09] | 0.02 |
| FX | -0.01 | 0.87/0.97 | [-0.09, 0.07] | 9.53 | -0.01 | 0.82/0.95 | [-0.10, 0.08] | 14.17 |
| FXST | -0.02 | 0.60/0.97 | [-0.10, 0.06] | 0 | -0.02 | 0.67/0.95 | [-0.10, 0.06] | 0 |
| GCC | -0.02 | 0.59/0.97 | [-0.10, 0.05] | 0 | -0.03 | 0.53/0.90 | [-0.10, 0.05] | 0 |
| IC | 0.00 | 0.96/0.97 | [-0.08, 0.07] | 0 | 0.00 | 0.90/0.95 | [-0.07, 0.08] | 0 |
| PCR | -0.03 | 0.37/0.97 | [-0.11, 0.04] | 0 | -0.03 | 0.52/0.90 | [-0.10, 0.05] | 0 |
| PLIC | 0.02 | 0.70/0.97 | [-0.06, 0.09] | 0.00 | 0.02 | 0.56/0.90 | [-0.06, 0.10] | 0 |
| PTR | 0.00 | 0.96/0.97 | [-0.08, 0.08] | 8.52 | 0.00 | 0.91/0.95 | [-0.09, 0.08] | 8.84 |
| RLIC | 0.01 | 0.89/0.97 | [-0.07, 0.08] | 0.00 | 0.01 | 0.85/0.95 | [-0.07, 0.09] | 0 |
| SCC | -0.06 | 0.15/0.86 | [-0.15, 0.02] | 14.30 | -0.04 | 0.33/0.90 | [-0.13, 0.04] | 13.44 |
| SCR | -0.02 | 0.63/0.97 | [-0.09, 0.06] | 0.00 | -0.02 | 0.53/0.90 | [-0.10, 0.05] | 0 |
| SFO | *-0.10* | *0.020/0.24* | *[-0.18, -0.02]* | *9.29* | *-0.11* | *0.0078/0.09* | *[-0.20, -0.03]* | *6.38* |
| SLF | 0.03 | 0.41/0.97 | [-0.04, 0.11] | 0.01 | 0.04 | 0.27/0.90 | [-0.04, 0.12] | 0.01 |
| SS | 0.00 | 0.97/0.97 | [-0.07, 0.08] | 0.01 | -0.01 | 0.73/0.95 | [-0.09, 0.07] | 0 |
| TAP | **-0.12** | **0.0021/0.050** | **[-0.19, -0.04]** | **0** | *-0.11* | *0.0044/0.094* | *[-0.19, -0.04]* | *0.02* |
| UNC | 0.01 | 0.77/0.97 | [-0.07, 0.09] | 0 | 0.00 | 0.96/0.96 | [-0.08, 0.08] | 0 |

Comparing trauma-exposed (n=200) to trauma un-exposed controls (n=93) from 6 sites, we found marginally lower FA in exposed controls in the tapetum, splenium of corpus callosum, and fornix/stria-terminalis (*d*=−0.41, *p*=0.014; *d*=−0.48, *p*=0.0042; *d*=−0.36, *p*=0.019, respectively), along with significantly higher MD and RD in the splenium (*d*=0.48, *p*=0.0017; *d*=0.56, *p*=0.00023, respectively) and marginally higher RD in the tapetum (*d*=0.32, *p*=0.036).

### Subgroups

We examined military vs. civilian cohorts, and male vs. female participants separately. All subgroups showed non-significant associations with PTSD, but there were marginal associations with tapetum FA in the military-only and male-only subgroups separately (see **Supplementary Note 3** and **Supplementary Figure 3** for more details). Results of group-by-sex and group-by-age interactions were not significant and are shown in **Supplementary Note 4** and **Supplementary Figure 4**.

### Additional Covariates

The role of potentially confounding variables on the association between PTSD and the tapetum was tested in several *post hoc* analyses focused on left, right, and bilateral tapetum FA (see **Supplementary Figure 2**). As these analyses were considered *post hoc* and limited to the tapetum, we used a test-wise significance threshold of *p*<0.05. Results generally remained significant across all models. Including dichotomous depression as a covariate (696 PTSD vs. 825 controls) resulted in lower left tapetum FA in the PTSD group (left *d*=−0.15, *p*=0.0090) and borderline lower bilateral tapetum FA (*d*=−0.11, *p*=0.090). Including AUD as a covariate (691 PTSD vs. 623 controls) resulted in lower bilateral and left tapetum FA in the PTSD group (bilateral *d*=−0.14, *p*=0.012; left *d*=−0.16, *p*=0.0061) and borderline lower right tapetum FA (*d*=−0.10, *p*=0.066). Including a binary TBI variable (849 PTSD vs. 1,016 controls) resulted in lower left tapetum FA (*d*=−0.12, *p*=0.015). Including a dichotomous psychotropic medication covariate (713 PTSD vs. 679 controls) resulted in lower bilateral and left tapetum FA in the PTSD group (bilateral *d*=−0.11, *p*=0.050; left *d*=−0.014, *p*=0.013). Including childhood trauma as a covariate (367 PTSD vs. 598 controls) did not yield any significant results, but neither did the analysis in the reduced sample, suggesting that the sample reduction impacted these results. To control for covariate- and cohort-dependent changes in sample size, each analysis was repeated in a smaller sample that corresponded to omitting the relevant covariate. The tapetum results remained consistent in nearly all reduced sample analyses – significant effects survived covariate adjustment and effects that disappeared (such as with childhood trauma) were also absent in the reduced sample. Thus, covariates had minimal impact beyond the reduction in sample size (see **Supplementary Note 5** and **Supplementary Figures 5–9**). A table showing how many participants at each site had information on these potentially confounding variables may be found in **Supplementary Table 3**.

### PTSD Severity

PTSD symptom severity in the PTSD group (measured by the CAPS-4, N=979 from 18 sites) was not associated with FA (**Figure 3**). Subgroup analyses yielded marginal associations between CAPS-4 score and tapetum FA in the military cohorts, but no other subgroup. Results were similar when potentially confounding variables were included, with no significant associations, although the tapetum was marginally significant when psychotropic medication use was included. Detailed analyses of PTSD severity and covariates within subgroups are in **Supplementary Note 5** and **Supplementary Figures 10** and **11**.

**Figure 3.**
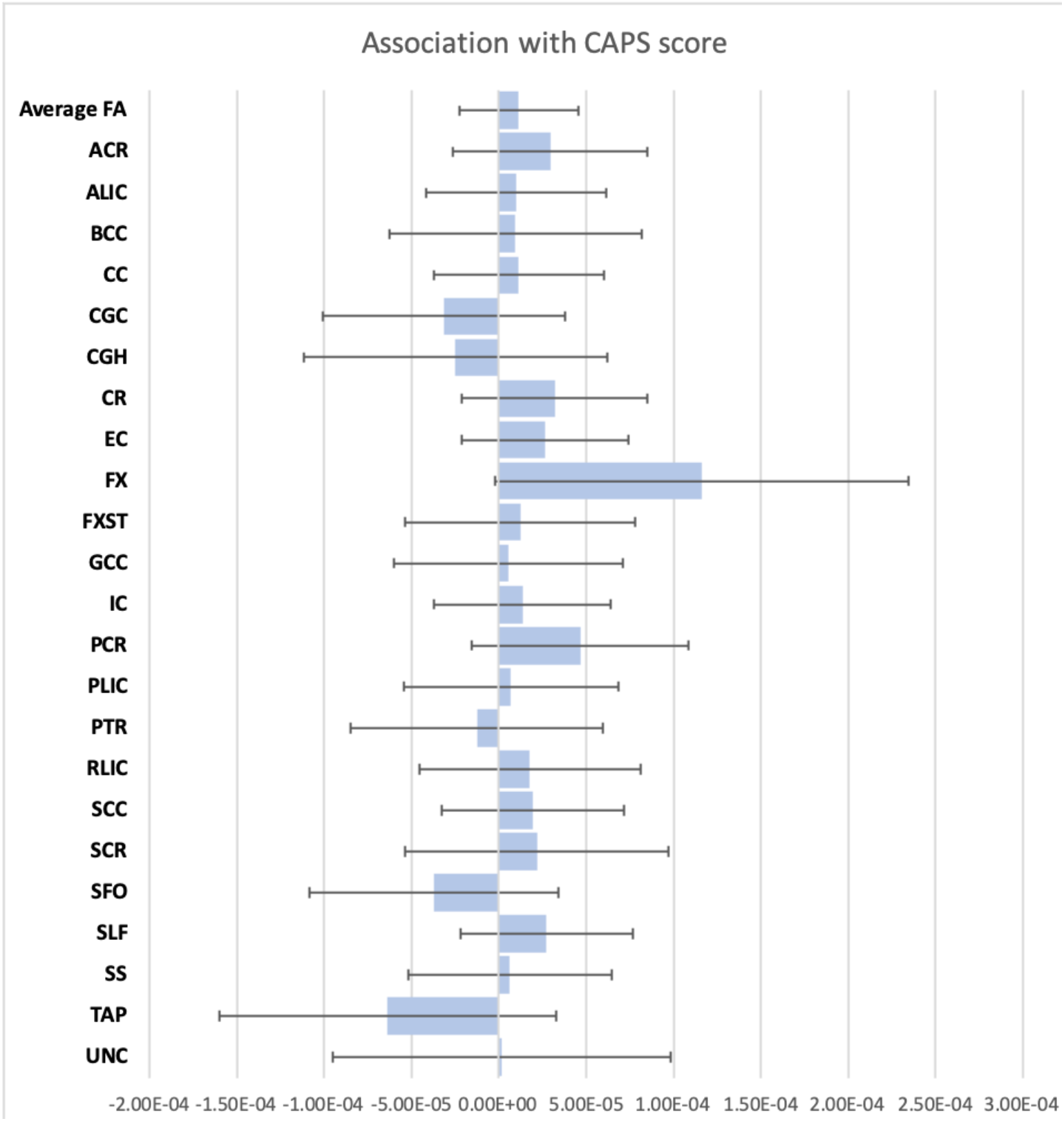
Linear association with CAPS-4 in the PTSD group. Meta-regression unstandardized β statistics are shown across all bilateral and midline ROIs and average FA, with bars indicating the 95% confidence interval. The ROI abbreviations are explained in **Supplementary Note 1**. Dark orange bars indicate significance (*p*<0.0021) and light orange bars indicates marginally significant results (0.05>*p*>0.0021).

## Discussion

We present DTI results from a multi-cohort study conducted by the PGC-ENIGMA PTSD consortium. In a meta-analysis of 3,049 participants from 28 sites, we found lower FA and higher RD in the tapetum among adults with PTSD (neuroanatomical figure – **Supplementary Figure 12**), which remained after accounting for several potentially confounding factors. The tapetum is a major tract within the corpus callosum that serves as a conduit between right and left hippocampus. Prior studies of white matter disruption in PTSD have found alterations in other hippocampal tracts, but were generally hindered by small sample sizes leading to inconsistent findings across studies. Our results add to the existing literature in identifying structural disruptions that compromise putative hippocampal functions, which are known to play a central role in PTSD symptomatology^48,49^. The tapetum is a segment of the corpus callosum that connects the temporal lobes, in particular the left and right hippocampus^50^. It is one of the last corpus callosum segments to develop and experiences rapid growth around age 14, which may make it vulnerable to the effects of trauma for a longer period of time^51^. Along with the inferior longitudinal fasciculus, cingulum, fornix, spino-limbic tracts (not studied here), and the anterior commissure, the tapetum is a dominant hippocampal pathway^50^. Structural and functional alterations in the hippocampus are frequently reported in PTSD, with smaller volumes^37^, decreased activation, and disrupted functional connectivity with the medial and lateral prefrontal cortices^52,53^. Here we report microstructural evidence that structural connectivity between the left and right hippocampus may also be disrupted in PTSD. While many studies have reported that PTSD is associated with alterations in the cingulum bundle^9–13,16,18,21,23–29,31,32,36^, which has a hippocampal component, the tapetum has not yet emerged for several possible reasons. Many prior studies took an ROI approach, which limited analyses to pre-determined regions that frequently omitted the tapetum, a small region often grouped with other tracts such as the splenium or posterior thalamic radiation. Critically, in 2013, an error was uncovered in the JHU atlas used as part of the TBSS pipeline, with the uncinate incorrectly identified as the inferior fronto-occipital fasciculus, and the tapetum incorrectly identified as the uncinate^54^. Thus, the tapetum was simply not examined in prior studies, with one very recent exception showing that tapetum abnormalities are associated with lower major depressive disorder remission^55^. Finally, the precise role of the tapetum in connecting the left and right hippocampus was only recently elucidated by mapping the subcortical connectome with exquisitely high-resolution mapping capable of discerning the intermingling of tapetum and other corpus callosum fibers. Super-resolution DTI conducted by the Chronic Diseases Connectome Project that acquired 1150-direction single-subject and 391-direction 94-subject data made this ultra-structural mapping possible^50^.

Childhood trauma is the greatest single risk factor for future vulnerability to PTSD^56^; numerous studies show significant alterations in brain structure and function in individuals who had experienced significant early life stress^46,57^. Some of these alterations likely contribute to a higher risk for psychopathology, but childhood trauma exposure did not explain the association between tapetum white matter disruption and PTSD that we report here. Depression is frequently comorbid with PTSD^58^ and is associated with disrupted white matter organization, although the affected tracts are broadly distributed^59^. Accounting for depression in group comparisons did not significantly alter our results, suggesting that tapetum white matter disruption is specific to PTSD. Particularly in military populations, which formed the majority of our sample, PTSD is often comorbid with traumatic brain injury (TBI)^60^. White matter is particularly vulnerable to TBI, which produces stretching and shearing of axons and altered neurometabolism^61^. Accounting for TBI also did not significantly change our results, indicating that TBI was associated with white matter damage generally, but not specifically within the tapetum. Psychotropic medications are another potential confound, given their neurotrophic and neuroprotective effects^62^. The result in the tapetum persisted after covarying for psychotropic medication, indicating that our findings are unlikely to be explained by medication. Lastly, PTSD can be comorbid with alcohol use disorders, which have a poorer clinical prognosis^63,64^. alcohol use disorders have been associated with significant changes in white matter organization^65,66^ but did not influence the present results.

We found a marginally significant association of PTSD with FA in the tapetum in male and military subgroups separately. Although results were non-significant in female or civilian subgroups, the effect size was slightly larger and in the same direction. The female and civilian subgroups were smaller and therefore the analyses had lower power than in male and military subgroups. Most prior dMRI studies in civilians report lower FA in PTSD^9,10,12–14,16,17,21–25,30–32,67^. Studies of military cohorts have been mixed, reporting higher FA^26–29,36^, lower FA^8,15,18–20^, and null results^33–35^. This discrepancy may be due to differences in age, chronicity, and type of trauma exposure, although military personnel often also experience civilian trauma. Combat-related PTSD is often comorbid with TBI, which is also associated with white matter disruption, constituting a potentially confounding factor for studies^68^.

In the absence of longitudinal data, our analysis cannot make causal inferences about the direction of the relationship between PTSD and tapetum white matter organization. Disrupted white matter of the tapetum may represent a vulnerability that predates the onset of PTSD, or a pathological response to trauma. In twins discordant for exposure to combat stress, the unexposed twins of combat veterans with PTSD have smaller hippocampal volume than the unexposed twins of combat veterans without PTSD^69^. Individuals with two risk alleles of the *FKBP5* gene have demonstrated lower cingulum FA above and beyond the association of cingulum FA with PTSD^11,67^. These studies suggest that heritable differences in brain structure may influence risk of developing PTSD. Evidence that alterations are caused by PTSD was observed in Israeli Defense Force recruits with reduced structural connectivity between the hippocampus and ventromedial prefrontal cortex, but only after exposure to military stress^70^. With the varying developmental trajectories of brain structure, function, and connectivity, along with the varying distribution of stress hormone receptors in the brain, the complex question of vulnerability vs. consequence will require prospective longitudinal neuroimaging studies.

Some evidence indicates that high FA is a marker of resilience to the effects of stress^71,72^. A putative marker of resilience is the ability to attenuate stress-induced increases in corticotropin-releasing hormone and glucocorticoids through an elaborate negative feedback system, and to modulate the expression of brain-derived neurotrophic factor (BDNF)^73,74^. BDNF has myriad functions including supporting neuronal differentiation, maturation, and survival^75,76^. In particular, hippocampal BDNF is implicated in the development of neural circuits that promote stress adaptations^73^. These stress adaptation circuits involve white matter in the fornix and other fronto-limbic connections^77^.

### Limitations

Our study has several limitations. One limitation of TBSS studies is the inability to fully attribute results to particular fiber bundles. Future studies may benefit by using tractography to more reliably identify the affected bundles, but this is difficult across so many varied sites. Second, not all participants classified as PTSD received clinician-administered interview (such as the CAPS) to confirm diagnoses. Third, we could not reliably measure chronicity across different cohorts. Other variables that we could not examine given the heterogeneity across sites include treatment effects, symptom clusters, trauma types, and lifetime as opposed to current PTSD diagnosis. Although we analyzed data from over 3,000 participants, we may have been underpowered to examine group-by-sex interactions, as 55% of our sample came from cohorts including only males or only females or samples that were >90% male. Diffusion metrics are not scanner invariant, and can vary even in scanners of the same model. For this reason, we were limited to a meta-analytic approach which may have lower power than mega-analysis. However, studies including both meta- and mega-analyses of brain volume in other ENIGMA groups have found minimal differences^78,79^.

Future studies should further investigate the tapetum using high spatial and angular resolution tractography to replicate our findings. Future and existing studies with more in-depth phenotyping than was possible here could examine how alterations in the tapetum vary with trauma type, chronicity, treatment, and whether they are associated with specific symptom clusters. The current study excluded pediatric cases, so additional research on white matter disruption in pediatric trauma and PTSD is warranted. Lastly, while we considered comorbidities as potential confounding variables, we did not examine their association with dMRI metrics. Future collaborations with the ENIGMA Brain Injury, MDD, and Addiction working groups will provide opportunities to separate general neuroimaging biomarkers of psychopathology and disorder-specific effects.

## Conclusions

Here we presented results from the PGC-ENIGMA PTSD working group, reporting poorer white matter organization in the tapetum in individuals currently suffering from PTSD. We present the largest DTI study in PTSD to date and the first to use harmonized image processing across sites, increasing our power to detect subtle effects. While future studies need to confirm the involvement of the tapetum specifically, our results add to the existing literature implicating the hippocampus as a primary area of disruption in PTSD.

## Acknowledgements

K99NS096116 to Dr. Emily Dennis; CIHR, CIMVHR; Dana Foundation (to Dr. Nitschke); the University of Wisconsin Institute for Clinical and Translational Research (to Dr. Emma Seppala); a National Science Foundation Graduate Research Fellowship (to Dr. Daniel Grupe); R01MH043454 and T32MH018931 (to Dr. Richard Davidson); and a core grant to the Waisman Center from the National Institute of Child Health and Human Development (P30HD003352); Defense and Veterans Brain Injury Centers, the U.S. Army Medical Research and Materiel Command (USAMRMC; W81XWH-13-2-0025) and the Chronic Effects of Neurotrauma Consortium (CENC; PT108802-SC104835); Department of Defense award number W81XWH-12-2-0012; ENIGMA was also supported in part by NIH U54EB020403 from the Big Data to Knowledge (BD2K) program, R56AG058854, R01MH116147, R01MH111671, and P41EB015922 to Dr. Paul Thompson and Dr. Neda Jahanshad; Department of Veterans Affairs via support for the National Center for PTSD, NIAAA via its support for (P50) Center for the Translational Neuroscience of Alcohol, and NCATS via its support of (CTSA) Yale Center for Clinical Investigation; DoD W81XWH-10-1-0925; Center for Brain and Behavior Research Pilot Grant; South Dakota Governor’s Research Center Grant; F32MH109274; Funding from the Bill & Melinda Gates Foundation; Funding from the SAMRC Unit on Risk & Resilience in Mental Disorders; German Research Foundation grant to J. K. Daniels (numbers DA 1222/4-1 and WA 1539/8-2); German Research Society (Deutsche Forschungsgemeinschaft, DFG; SFB/TRR 58: C06, C07); I01-CX000715 & I01-CX001542; K01MH118428; K23MH090366; K2CX001772, Clinical Science Research and Development Service, VA Office of Research and Development; MH098212; MH071537; M01RR00039; UL1TR000454; HD071982; HD085850; MH101380; NARSAD 27040; NARSAD Young Investigator, K01MH109836, Young Investigator Grant, Korean Scientists and Engineers Association; NHMRC Program Grant # 1073041; R01AG050595, R01AG022381; R01AG058822; R01EB015611; R01MH105355; R01MH105355, R01DA035484; R01MH111671, R01MH117601, R01AG059874, MJFF 14848; R01MH111671; VISN6 MIRECC; R21MH112956, Anonymous Women’s Health Fund, Kasparian Fund, Trauma Scholars Fund, Barlow Family Fund; South African Medical Research Council for the “Shared Roots” Flagship Project, Grant no. MRC-RFA-IFSP-01-2013/SHARED ROOTS” through funding received from the South African National Treasury under its Economic Competitiveness and Support Package. The work by Dr. Leigh van Heuvel reported herein was made possible through funding by the South African Medical Research Council through its Division of Research Capacity Development under the SAMRC Clinician Researcher (M.D PHD) Scholarship Programme from funding received from the South African National Treasury; South African Research Chairs Initiative in Posttraumatic Stress Disorder through the Department of Science and Technology and the National Research Foundation.; the National Natural Science Foundation of China (No. 31271099, 31471004), the Key Research Program of the Chinese Academy of Sciences (No. ZDRW-XH-2019-4); The study was supported by ZonMw, the Netherlands organization for Health Research and Development (40-00812-98-10041), and by a grant from the Academic Medical Center Research Council (110614) both awarded to Dr. Miranda Olff; Translational Research Center for TBI and Stress Disorders (TRACTS), a VA Rehabilitation Research and Development (RR&D) Traumatic Brain Injury Center of Excellence (B9254-C) at VA Boston Healthcare System; VA CSR&D 1IK2CX001680; VISN17 Center of Excellence Pilot funding; VA CSR&D 822-MR-18176 and Senior Career Scientist Award; VA National Center for PTSD; VA RR&D 1IK2RX000709; VA RR&D 1K1RX002325; 1K2RX002922; VA RR&D I01RX000622; CDMRP W81XWH-08–2–0038; W81XWH08-2-0159 to Dr. Murray Stein from the US Department of Defense. The views reflected here are strictly those of the authors and do not constitute endorsement by any of the funding sources listed here.

## Supplementary Information

**Supplementary Note 1.** Further detail of ENIGMA-DTI protocols

**Supplementary Note 2.** Details on depression inventories.

**Supplementary Note 3.** Subgroup analyses.

**Supplementary Note 4.** Interaction analyses.

**Supplementary Note 5.** Group and severity analyses covarying for childhood trauma, depression, alcohol use disorders, traumatic brain injury, or psychotropic medication use.

**Supplementary Table 1.** Clinical details and inclusion and exclusion criteria for each of the cohorts included.

| Site | Inclusion criteria | Exclusion criteria |
| --- | --- | --- |
| **ADNI-DoD** | PTSD: Subjects must be Veterans of the Vietnam War, 50-90 years of age. Subjects who meet the SCID-I (for DSM-IV-TR) criteria for current/chronic PTSD (identified by records and verified by our telephone assessments). In addition to meeting DSM-IV-TR criteria for current/chronic PTSD, subjects must have a minimum current CAPS score of 50 as determined by telephone assessment. The PTSD symptoms contributing to the PTSD Diagnosis and Current CAPS score must be related to a Vietnam War related trauma. Must live within 150 miles of the closest ADNI clinic in subject's area. Control: Subjects must be Veterans of the Vietnam War, 50-90 years of age. Comparable in age, gender, and education with TBI and PTSD groups May be receiving VA disability payments for something other than TBI or PTSD – or no disability at all. Must live within 150 miles of the closest ADNI clinic in subject's area | PTSD: Mild Cognitive Impairment/Dementia Documented or self report history of mild/moderate severe TBI Any history of head trauma associated with injury onset cognitive complaints, or Loss of consciousness for >5minutes. Control: MCI/Dementia Presence of PTSD by SCID-I for DSM-IV-TR criteria, or a CAPS score of >30 (Both current and/or a history of PTSD will be excluded). Documented or self report history of mild/moderate severe TBI Any history of head trauma associated with injury onset cognitive complaints, or Loss of Consciousness for >5 minutes History of PTSD or current PTSD Exclusionary criteria applied to TBI/PTSD will be applied to controls. ALL: MCI/dementia History of psychosis or bipolar affective disorder; History of alcohol or substance abuse/dependence within the past 5 years (by DSM IV – TR criteria); MRI-related exclusions: aneurysm clips, metal implants that are determined to be unsafe for MRI; and/or claustrophobia; Contraindications for lumbar puncture, PET scan, or other procedures in this study; Any major medical condition must be stable for at least 4 months prior to enrollment. These include but are not limited to clinically significant hepatic, renal, pulmonary, metabolic or endocrine disease, cancer, HIV infection and AIDS, as well as cardiovascular disease. Seizure disorder or any systemic illness affecting brain function during the past 5 years will be exclusionary Clinical evidence of stroke. Have a history of relevant severe drug allergy or hypersensitivity. Subjects with current clinically significant unstable medical comorbidities, as indicated by history or physical exam, that pose a potential safety risk to the subject. |
| **Beijing** | (1) household was used as the basic sample unit, and the household member whose birthday was closest to the date of investigation was first selected for participation, and if the individual was unavailable, the household member whose birthday was the next closest was selected; (2) 38–62 years, and experienced the disaster personally; (3) right-handed. | Individuals with (1) mental retardation and any major psychosis (e.g., schizophrenia and organic mental disorders); (2) drug or alcohol abuse; (3) history of head trauma or surgery;(4) a metallic embedded object in body; (5) claustrophobia; (6) exposure to other trauma events from time of the disaster to the time of the study. |
| **Booster** | All: 18-65 years of age, police officers, eligible for MRI. PTSD: current PTSD diagnosis, with CAPS ≥ 45. Controls: exposure to at least one traumatic event (according to DSM-IV A1 criterion), with CAPS < 15 | All: history of neurological disorders, any severe or chronic systemic disease or unstable medical condition (including endocrinological disorders), use of psychotropic medications. Females: pregnancy or breastfeeding. PTSD: current psychotic disorder, substance-related disorder, severe personality disorder, severe major depressive disorder (MDD) (i.e., involving high suicidal risk and/or psychotic symptoms) or current suicidal risk. Controls: any current Axis-1 disorder and lifetime history of PTSD or MDD |
| **Columbia** | All: Males or females between the ages of 18 and 60, able to give consent, fluent in English; PTSD: Experience of a traumatic event or events in childhood and/or adulthood; current DSM-V Criterion A for PTSD | 1. Prior or current Axis I psychiatric diagnosis of schizophrenia, psychotic disorder, bipolar disorder, dementia. SCID and clinical evaluation 2. Depression score of > 25 on the Hamilton Rating Scale for Depression (HAM-D-17-item); significant depression and /or depression related impairment that is judged to warrant pharmacotherapy or combined medication and psychotherapy. Clinical interview, HAM-D. 3. Individuals at risk for suicide based on history and current mental state. Psychiatric history; mental status exam by screening psychiatrist; score > 2 on item 3 of Hamilton Depression Scale 4. History of substance/alcohol dependence within the past six months, or abuse within past two months History, urine toxicology 5. Any psychotropic medications including: antipsychotic, antidepressant, mood stabilizer, or stimulant medications in the last 4 weeks prior to the study (6 weeks for fluoxetine). Standing daily dosing of benzodiazepine class of medication in the 2 weeks prior to the study (as needed use of benzodiazepines is not an exclusion, but must be clinically judged to tolerate no benzodiazepines for the 72-hour period before each of the fMRI days). Triptan anti-migraine medications. Other medications that may interfere with fear circuitry and fear memory such as blood-brain-barrier penetrating β-blockers. Clinical Interview regarding current medication treatment. History. 7. Pregnancy, or plans to become pregnant during the period of the study. Urine β-HCG, interview 8. Paramagnetic metallic implants or devices contraindicating magnetic resonance imaging or any other non-removable paramagnetic metal in the body. History and MRI-Screening Questionnaire 9. Medical illness that could interfere with assessment of diagnosis, or |
|  |  | biological measures (SCR, fMRI), including organic brain impairment from stroke, CNS tumor, or demyelinating disease; and renal, thyroid, hematologic or hepatic impairment Medical chart review, clinical interview, physical exam, blood chemistry 10. Current unstable or untreated medical illness; or resting SBP>140 or DBP>90; or HR<60 or HR>100 History and physical exam 11. Any condition that would exclude MRI exam (e.g. pacemaker, paramagnetic metallic prosthesis, surgical clips, shrapnel, necessity for constant medicinal patch, some tattoos) Clinical interview, physical exam, MRI screening Questionnaire 12. Significant claustrophobia that would preclude ability to remain calm within the MRI scanner Interview, history; Controls: 1. Current symptomatic major Axis I psychiatric diagnosis, e.g., major depressive disorder, psychotic disorder, bipolar disorder, obsessive compulsive disorder (OCD), PTSD, panic disorder, agoraphobia, or eating disorder. Psychiatric Interview and SCID 2. Lifetime history of any DSM-5 Axis I disorder Psychiatric Interview and SCID 3. History of trauma exposure that fulfills DSM-5 PTSD criteria A1 History, Life Event Checklist (LEC), SCID 4. Depression score greater than 7 on the Hamilton Rating Scale for Depression (HAM-D-17-item) HAM-D 5. Lifetime history of substance/alcohol dependence or abuse History, urine toxicology 6. Pregnancy, or plans to become pregnant during the period of the study. Interview, Urine β-HCG for women of childbearing potential on each day of fMRI/emotional learning task. 7. Medical illness that could interfere with assessment of response or biological measures (SCR, fMRI), including organic brain impairment from stroke, CNS tumor, or demyelinating disease; or renal, thyroid, hematologic or hepatic impairment Medical chart review, clinical interview, physical exam, blood chemistry 8. Paramagnetic metallic implants or devices contraindicating magnetic resonance imaging or any other non-removable paramagnetic metal in the body. History and MRI-Screening Questionnaire 9. Medical illness that could interfere with assessment of diagnosis or biological measures (SCR, fMRI), including organic brain impairment from stroke, CNS tumor, or demyelinating disease; and renal, thyroid, hematologic or hepatic impairment. Medical review of systems, clinical interview, blood pressure and pulse assessment 10. Current unstable or untreated medical illness or resting SBP>140 or DBP>90; or HR<59 or HR>100 History and physical exam 11. Any condition that would exclude MRI exam (e.g. pacemaker, paramagnetic metallic prosthesis, surgical clips, shrapnel, necessity for constant medicinal patch, some tattoos) Clinical interview, physical exam, MRI screening Questionnaire 12. Significant claustrophobia that would preclude ability to remain calm within the MRI scanner |
| Duke (all) | All: 18-65, OEF/OIF veterans, fluent in English, free of implanted metal objects or metal shards in eyes, antidepressant, sleep, and anti-anxiety medication permitted | All: Axis I other than PTSD or MDD, current substance abuse or lifetime substance dependence (other than nicotine), high risk for suicide, claustrophobia, neurological disorders, learning disability or developmental delay, major medical conditions |
| Grady Trauma Project | All: 18-65 years of age, English-speaking, endorsed at least 1 criterion A trauma | All: Current psychotic symptoms or bipolar disorder, current substance or alcohol dependence, history of head trauma, taking any psychoactive medication, current illegal drug use (verified with urine drug screen within 24 hours of scan) |
| Groningen | 20-60 years, women with current PTSD diagnosis, civilian, sufficient proficiency in German, MRI compatible | Neurologic disorders, history of substance abuse or dependence within the last 6 months, history of head injury, cerebral incidental findings verified by a neuroradiologist after the MR scan, intake of benzodiazepines, tricyclic antidepressants, anticonvulsants, primary borderline personality disorder, current diagnosis of Axis I disorder |
| INTRuST | For complete inclusion and exclusion criteria, see <sup>10</sup> |  |
| iSCORE | PTSD: Age 25-48, English fluency, individuals were screened for a current PTSD diagnosis using the Weather's et al criterion for the PTSD checklist (PCL), a score of 44 or higher. PTSD symptoms were further required to be combat related and participants had to have returned from combat related deployment within three to 36 months of study enrollment. | Any participants with conditions preventing MRI procedures (i.e. claustrophobia, shrapnel, pregnancy), neurologic conditions (e.g., seizures, psychosis, etc.), history of TBI exceeding mild severity or spinal cord injury, or current narcotic medicine use were excluded from the study. |
| Lawson | Primary diagnosis of PTSD | Psychotic disorder, bipolar disorder, traumatic brain injury, narcotic use, active substance use disorder within 3 months of study entry |
| McLean | History of childhood maltreatment; Legal and mental competency of the patient; Female; All ethnic backgrounds; Age between 18 and 60; Fluent English speakers; Normal or Corrected Vision | Delirium secondary to medical illness; History of neurological conditions that may cause significant psychiatric symptomatology (e.g., dementia); Any contraindication to MR scans, including claustrophobia, pregnancy, metal implants, etc.; Current alcohol or substance use disorder (within the last month); A history of schizophrenia or other psychotic disorder; History of head injury or loss of consciousness for longer than 5 min (including concussion); Positive pregnancy test |
| Münster | IPV trauma, normal or corrected-to-normal vision, right-handed | na |
| New South Wales | 18-60 years | Neurological disorders, traumatic brain injury, psychosis |
| South Dakota | Study 1: OIF/ OEF/OND (Operation Iraqi Freedom / Operation Enduring Freedom / Operation New Dawn) veterans | Study 1: (a) Current or previous seizure history; b) current crisis-related issues such as serious self-injurious behavior, psychosis, or substance dependence (excluding alcohol dependence); (c) report of traumatic brain injury using the Traumatic Brain Injury Checklist (Hoge et al., 2008); and (d) contraindications to fMRI (metal objects in body, claustrophobia) |
| Stellenbosch (both) | Adult patients aged ≥18 years with a diagnosis of current PTSD. Able to read and understand the Informed Consent documents and be able to read and write in English or Afrikaans. | Any other major psychiatric disorder (e.g. severe mood and psychotic disorders) or any neurological disorder, and significant head injury, any alcohol or drug use disorder within the past 6 months. Pregnancy. |
| U Sydney | 18-65 years of age, endorsed at least 1 criterion A trauma, English-speaking | History of neurological illness (Huntington's, Parkinson's, dementia, MS, etc). History of Seizure Disorders, unrelated to head injury(ies). Current diagnosis of schizophrenia spectrum or other psychotic disorders (not related to PTSD). Current diagnosis of bipolar or related disorders (not related to PTSD). Current active homicidal and/or suicidal ideation with intent requiring crisis intervention. Cognitive disorder due to general medical condition other than TBI. Unstable psychological diagnosis that would interfere with accurate data collection, determined by consensus of at least two doctorate-level psychologists. Also, MRI contra-indications for MRI including metallic implants or foreign objects deemed unsafe by the MRI technician (such as but not limited to pace-maker, shrapnel, metallic screws). Surgery in the past 2 months except as approved by the MRI technician (such as but not limited to dental work, colonoscopy). Weight exceeding the capacity of the scanner table. |
| UMC Utrecht | All: 18-60 years of age, eligible for MRI. PTSD: current PTSD diagnosis, with CAPS ≥ 45, military deployment >4 months. Trauma controls: exposure to at least one traumatic event (according to DSM-IV A1 criterion), with CAPS < 15, no current psychiatric disorder, military deployment >4 months; healthy controls: no current psychiatric disorder according to DSM-IV. | All: history of neurological disorders, any severe or chronic disorder; alcohol or drug abuse and/or dependence during course of the study. |
| UW-Madison | Age range of 18-50; Capable of giving informed consent; Fluent in English; Exposure to one or more life-threatening war zone trauma events per the Combat Experiences Scale and documented by DD-214, Combat Action Ribbon (Marines), Combat Infantry Badge (Army), or other clear evidence of war zone trauma exposure in Iraq or Afghanistan since 2001; Pharmacological or psychotherapeutic treatment | Weight of 352 pounds or over (due to constraints of MRI scanner); Women of childbearing potential with positive pregnancy test, looking to conceive during the research timeline, or who are breastfeeding; Metallic implants such as prostheses or aneurysm clip, or electronic implants such as cardiac pacemakers; Neurological or serious medical condition that may contraindicate MRI or that may overlap with physiological substrates of psychiatric conditions; History of seizures or seizure disorder; Moderate or severe traumatic brain injury (over 20 minutes unconscious); Current active substance dependence or dependence within 3 months (other than nicotine); Meets DSM-IV criteria for bipolar disorder, |
|  | stable for at least 8 weeks prior to beginning of study, with no intent to begin a new course of treatment during the study period | schizophrenia, schizoaffective disorder, psychotic disorder NOS, delirium, or any DSM-IV cognitive disorder.; Substance dependence disorder within 3 months or any current substance dependence; Severe psychiatric instability or severe situational life crises, including evidence of being actively suicidal or homicidal, or any behavior that poses an immediate danger to patient or others.; Participants with extensive experience in yoga or meditation; Current use of benzodiazepines and beta-blockers |
| VA Boston | 1. Veteran of OEF/OIF/OND (deployed at least one time to either Afghanistan or Iraq) or 2. Active duty Service Member (SM) not yet deployed to OEF/OIF/OND. Age 18-65 years. | History of neurological illness (Huntington's, Parkinson's, dementia, MS, etc). History of Seizure Disorders, unrelated to head injury(ies). Current diagnosis of schizophrenia spectrum or other psychotic disorders (not related to PTSD). Current diagnosis of bipolar or related disorders (not related to PTSD). Current active homicidal and/or suicidal ideation with intent requiring crisis intervention. Cognitive disorder due to general medical condition other than TBI. Unstable psychological diagnosis that would interfere with accurate data collection, determined by consensus of at least two doctorate-level psychologists. Also, MRI contra-indications for MRI including metallic implants or foreign objects deemed unsafe by the MRI technician (such as but not limited to pace-maker, shrapnel, metallic screws). Surgery in the past 2 months except as approved by the MRI technician (such as but not limited to dental work, colonoscopy). Weight exceeding the capacity of the scanner table. |
| VA Houston | All: Age 18-50 years, Right-handed (as determined by the Edinburgh Handedness Inventory by a score $\geq 40$), Able to safely and comfortably undergo MRI (e.g. no implanted metal, no embedded shrapnel, claustrophobia, etc.), participants are required to have been previously deployed in OEF/OIF/OND. PTSD: Meets DSM-V criteria for PTSD, meets DSM-V criteria for depression. Control: Uninjured, or sustaining a non-cranial injury without history of blast exposure, Post-injury interval $\geq 3$ months or more since most recent injury (for the participants with non-cranial injuries) | All: Not fluent in English, Ever having been formally diagnosed with dyslexia or other learning disabilities, Left-handed (as determined by the Edinburgh Handedness Inventory by a score $< 40$), Post-deployment TBI requiring hospitalization (i.e., not just treated and released directly from Emergency Center ), Contraindications to undergoing MR imaging (e.g., metal implants, orthodontia, shrapnel, positive urine pregnancy screen, claustrophobia, etc.), Pre- or post-deployment neurologic disorder associated with cerebral dysfunction and/or cognitive deficit (e.g., mental retardation, HIV/AIDS, dementias, etc.), Pre-deployment major psychiatric disorder associated with cerebral dysfunction and/or cognitive deficit (e.g., schizophrenia and other psychotic disorders, bipolar disorder), as determined by the Structured Clinical Interview for the DSM-V conducted by ROVER-WISER clinicians (or the most current version of the MINI for Veteran Controls—see criteria for Controls below), Current active psychosis, Any history of intracranial surgery, Medical conditions associated with structural/functional compromise on MRI or cognitive decrements; PTSD: Meets criteria for TBI group; control: 1. Meets MINI criteria for any substance use disorders (SUDs) other than nicotine (tobacco products), Meets MINI criteria for current depressive episode, Meets MINI criteria for PTSD, Meets MINI criteria for other major psychiatric disorder associated with cerebral dysfunction and/or cognitive deficit (e.g., schizophrenia and other psychotic disorders, bipolar disorder), Meets criteria for history of TBI of any severity |
| VA Minneapolis (both) | Participants were veterans of Operation Enduring Freedom and/or Operation Iraqi Freedom, age 22-60, who had been exposed to combat during their deployment(s). | Participants were excluded from the study if they met criteria for 1) a current substance-induced psychotic disorder or psychotic disorder due to a general medical condition (other than TBI), 2) current DSM-IV substance abuse or dependence other than alcohol, caffeine, or nicotine, 3) a moderate or severe traumatic brain injury from either impact or blast, 4) a neurologic condition other than TBI, 5) a current unstable medical condition that would likely affect brain function (e.g., uncontrolled diabetes), or 6) significant imminent risk of suicidal or homicidal behavior. |
| VA Waco | Study 1: Veterans ages 18-60; Study 2: Veterans age 18-60 with a clinical diagnosis of TBI in their VA medical record | Study 1: Seizure disorder, dementia, or MRI safety concerns; Study 2: 1) diagnosis of schizophrenia, schizoaffective disorder, bipolar disorder type I, severe substance use disorder, or a high risk of suicide (via the MINI); 2) absence of qEEG more than 2SDs outside of population means of healthy age-matched controls (in this study, EEG neurofeedback was conducted after scanning); and 3) MRI safety concerns. |
| VETSA | Participants were recruited from the Vietnam Era Twin Registry aka VETR (by registry definition, then, both brothers had been in the US military at some point between 1965 and 1975; not "VA"). Participants had to be between 50 to 59 years old when recruited; both brothers needed to agree to participate. | For MRI component, participants needed to meet safety criteria. |
| Yale/NCPTSD | Participants ranged in age from 21 to 60 and had been deployed on one or more combat tours. | Individuals were excluded from the study if they met any of the following criteria: a diagnosis of bipolar disorder or psychotic disorder, as assessed by the SCID-IV (First, Spitzer, Gibbon, & Williams, 2002); current benzodiazepine use; a history of ADHD, learning disorder, moderate or severe traumatic brain injury (TBI), brain tumor, epilepsy, or a neurological disorder; current inpatient status; or an MRI contraindication. |

**Supplementary Table 2.** DTI acquisition parameters for each of the cohorts included.

| Site | Scanner | Field strength | Voxel size (mm) | Gradient directions and b-value (mm/s<sup>2</sup>) | b0 scans |
| --- | --- | --- | --- | --- | --- |
| ADNI-DoD | 1. GE Discovery MR750, 2. GE Discovery MR750w, 3. GE Signa HDxt, 4. Siemens Tim Trio | 3T | 2x2x2 | 41 at b=1000 | 5 |
| Beijing | Philips | 1.5T | 1.71875x1.71875x2 | 16 at b=800 | 1 |
| Booster | Philips Achieva | 3T | 2x2x2 | 32 at b=1000 | 1 |
| Columbia | GE Discovery MR750 | 3T | 1.875x1.875x2.5 | 64 at b=1000 | 5 |
| Duke-1 | GE Discovery MR750 | 3T | 2x2x2 | 64 at b=900 | 5 |
| Duke-2 | Philips Ingenia | 3T | 2x2x2 | 32 at b=800 | 2 |
| Duke-3 | GE Discovery MR750 | 3T | 2x2x2 | 55 at b=1000 | 1 |
| Grady Trauma Project | Siemens Trio | 3T | 2x2x2 | 60 at b=1000 | 1 |
| Groningen | Siemens Trio | 3T | 2.3x2.3x2.3 | 64 at b=1000 | 1 |
| INTRuST | Philips | 3T | 2x2x2 | 87 at b=900 | 7 |
| iSCORE | Siemens Verio Syngo | 3T | 2x2x2 | 64 at b=1000 | 1 |
| Lawson | Siemens | 3T | 2x2x2 | 30 at b=1000 | 1 |
| McLean | Siemens Tim Trio | 3T | 2x2x2 | 48 at b=700 | 7 |
| Münster | Siemens Magnetom Prisma Fit Syngo MR D13D | 3T | 1.8x1.8x1.8 | 30 at b=1000 | 12 |
| New South Wales | GE Signa HDx | 3T | 1.72x1.72x2.5 | 42 at b=1250 | 4 |
| South Dakota | Siemens Skyra | 3T | 2x2x2 | 30 at b=1000 | 1 |
| Stellenbosch-1 | Siemens Skyra | 3T | 1.92x1.92x2.4 | 45 at b=1000 | 3 |
| Stellenbosch-2 | Siemens Allegra | 3T | 2x2x2.5 | 30 at b=1000 | 5 |
| U Sydney | GE Discovery MR750 | 3T | 2x2x2 | 69 at b=1000 | 2 |
| UMC Utrecht | Philips Achieva | 3T | 1.875x1.875x2 | 30 at b=1000 | 1 |
| UW Madison | GE Discovery MR750 | 3T | 2x2x2 | 66 at b=1200 | 4 |
| VA Boston | Siemens | 3T | 2x2x2 | 60 at b=700 | 10 |
| VA Houston | Siemens Trio | 3T | 2.67x2.67x3.25 | 30 at b=1000 | 1 |
| VA Minneapolis-1 | Siemens Tim Trio | 3T | 2x2x2 | 30 at b=800 | 10 |
| VA Minneapolis-2 | Siemens Tim Trio | 3T | 2x2x2 | 128 at b=1500 | 17 |
| VA Waco | Philips Achieva | 3T | 1.75x1.75x2 | 32 at b=800 | 1 |
| VETSA | 1. GE Discovery MR750, 2. Siemens Tim Trio | 3T | 2.5x2.5x2.5 | 1. 51 at b=1000, 2. 30 at b=1000 | 1. 2, 2. 1 |
| Yale/NCPTSD | Siemens Tim Trio | 3T | 1.7x1.7x3 | 128 at b=1000 | 1 |

**Supplementary Table 3.**
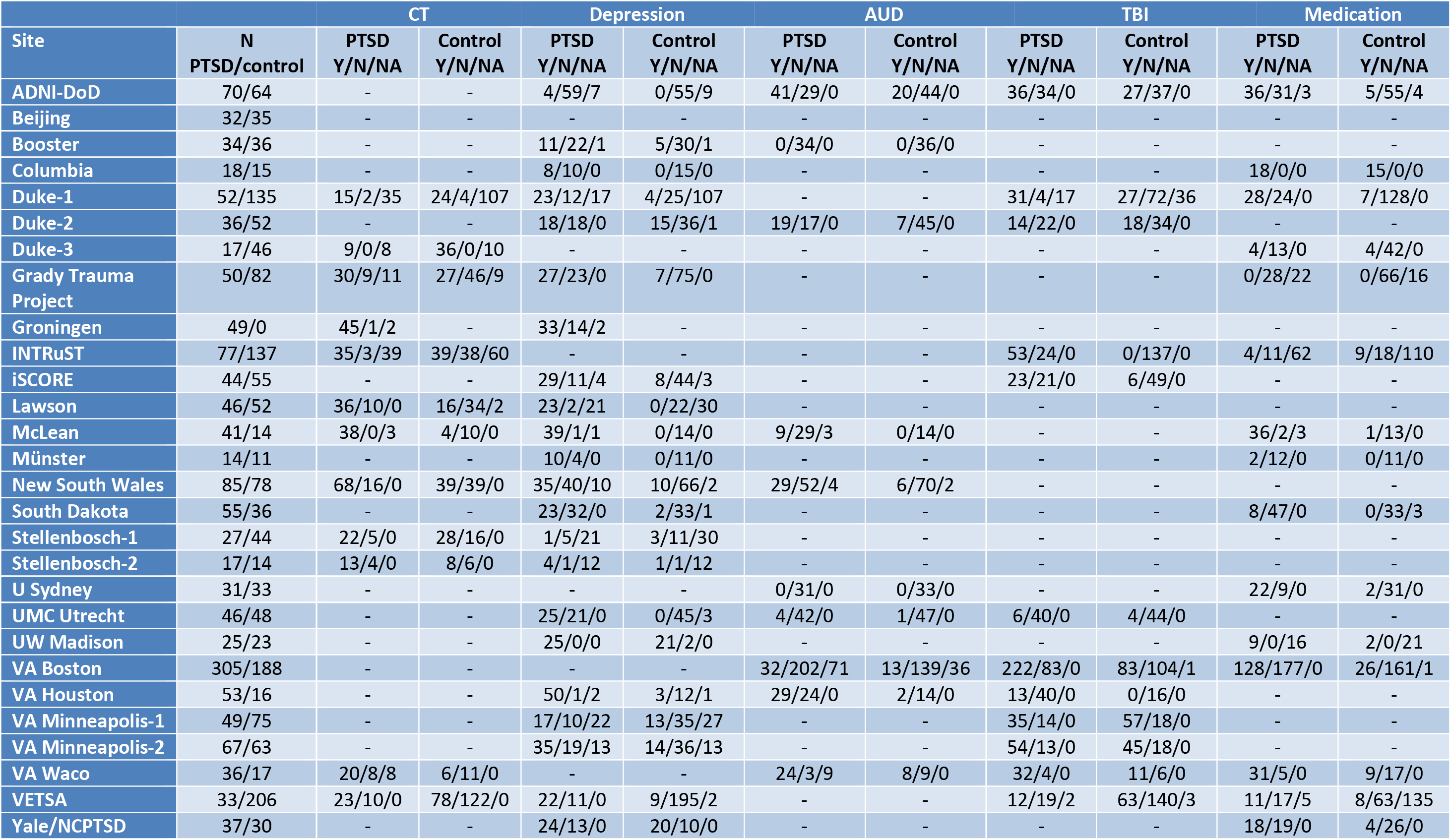
Number of participants per site with information on the potentially confounding covariates: childhood trauma (CT – binarized here), depression, alcohol use disorder (AUD – binarized here), traumatic brain injury (TBI), and psychotropic medications. Counts are given per site, per variable, per group for yes/no/NA (missing), with “-” for information that was not collected.

**Supplementary Figure 1.**
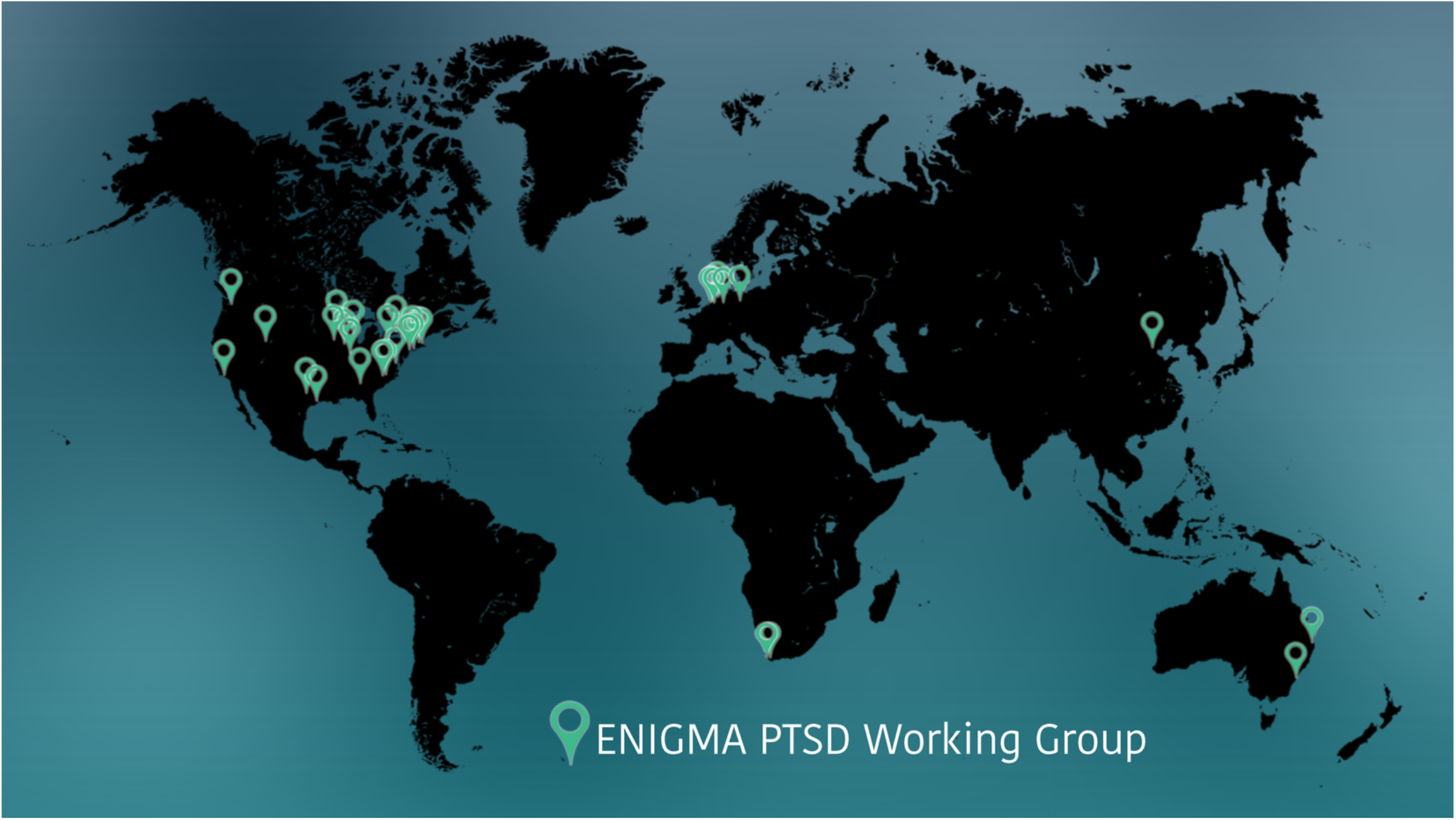
Map of cohorts included.

**Supplementary Figure 2.**
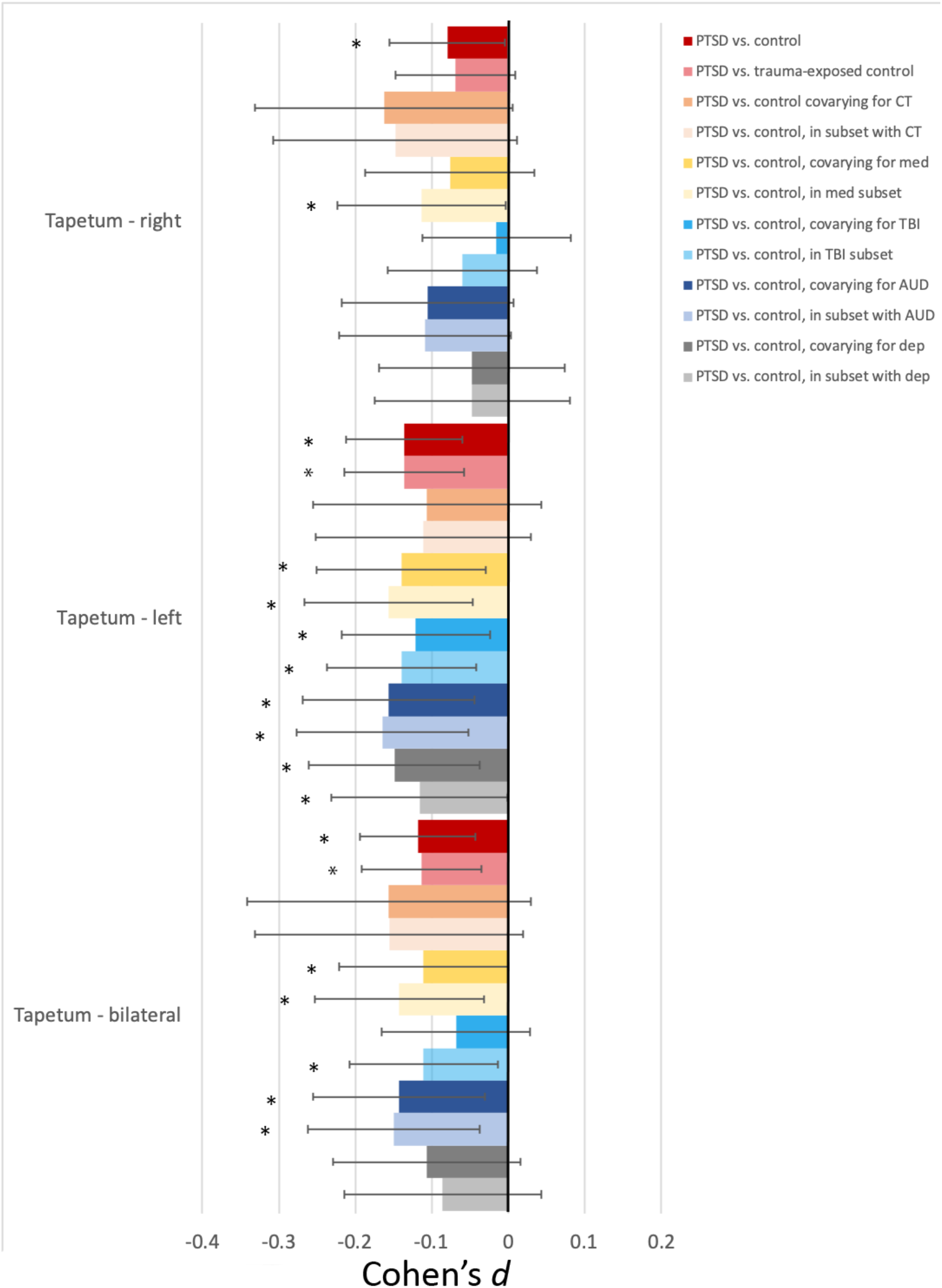
*Post-hoc* examination of tapetum result. Shown are the Cohen’s *d* for the left, right, and bilateral tapetum FA with 95% CI. Colors correspond to the models tested, as shown in the legend. Red/pink=PTSD vs. control/PTSD vs. trauma-exposed controls, orange/light orange=PTSD vs. control covarying for childhood trauma (CT)/in subset with CT without CT in model, yellow/light yellow= PTSD vs. control covarying for psychotropic medications (med)/in subset with med without med in model, teal/light teal= PTSD vs. control covarying for traumatic brain injury (TBI)/in subset with TBI without TBI in model, navy/light navy= PTSD vs. control covarying for alcohol use disorders (AUD)/in subset with AUD without AUD in model, gray/light gray= PTSD vs. control covarying for depression (dep)/in subset with dep without dep in model.

**Supplementary Figure 3.**
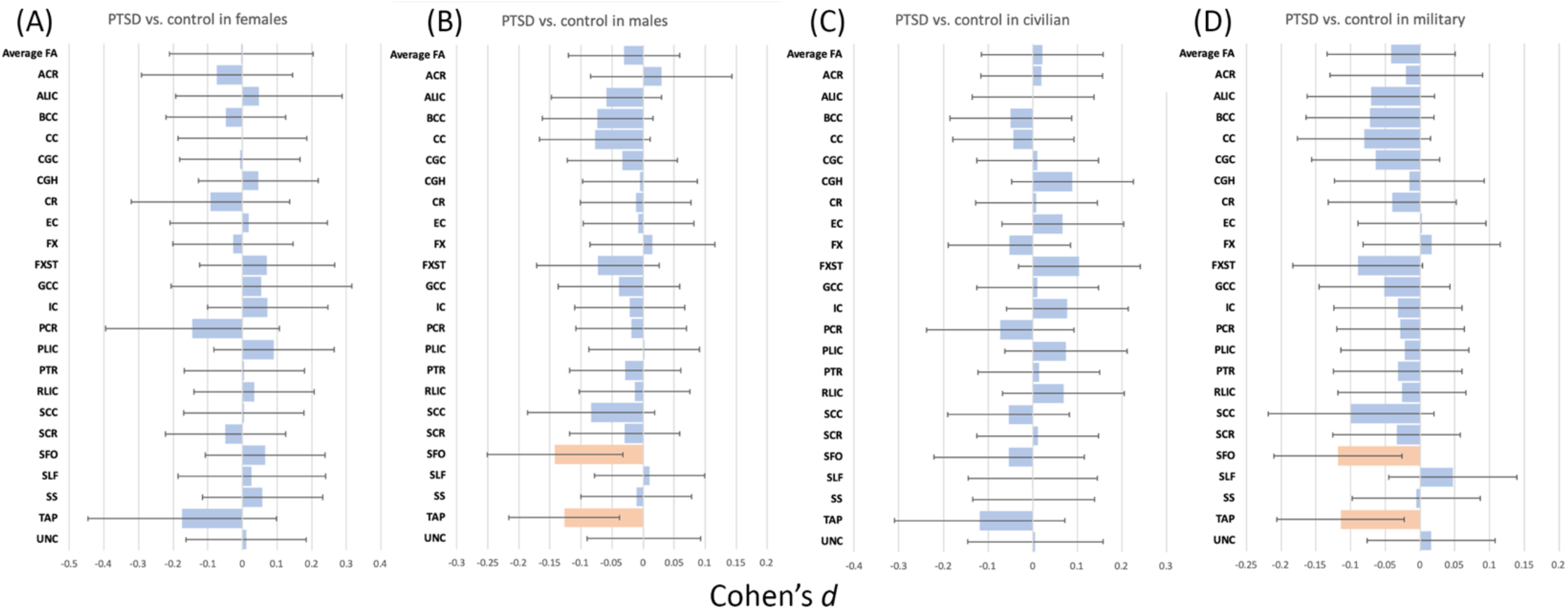
Subgroups: (a) PTSD vs. control effects in females only, (b) PTSD vs. control effects in males only, (c) PTSD vs. control effects in civilians only, and (d) PTSD vs. control effects in military only. Shown are Cohen’s *d* for 23 ROIs and average FA. Dark orange bars indicate significance (*p*<0.0021) and light orange bars indicate marginally significant results (0.05>*p*>0.0021). Error bars are 95% CI.

**Supplementary Figure 4.**
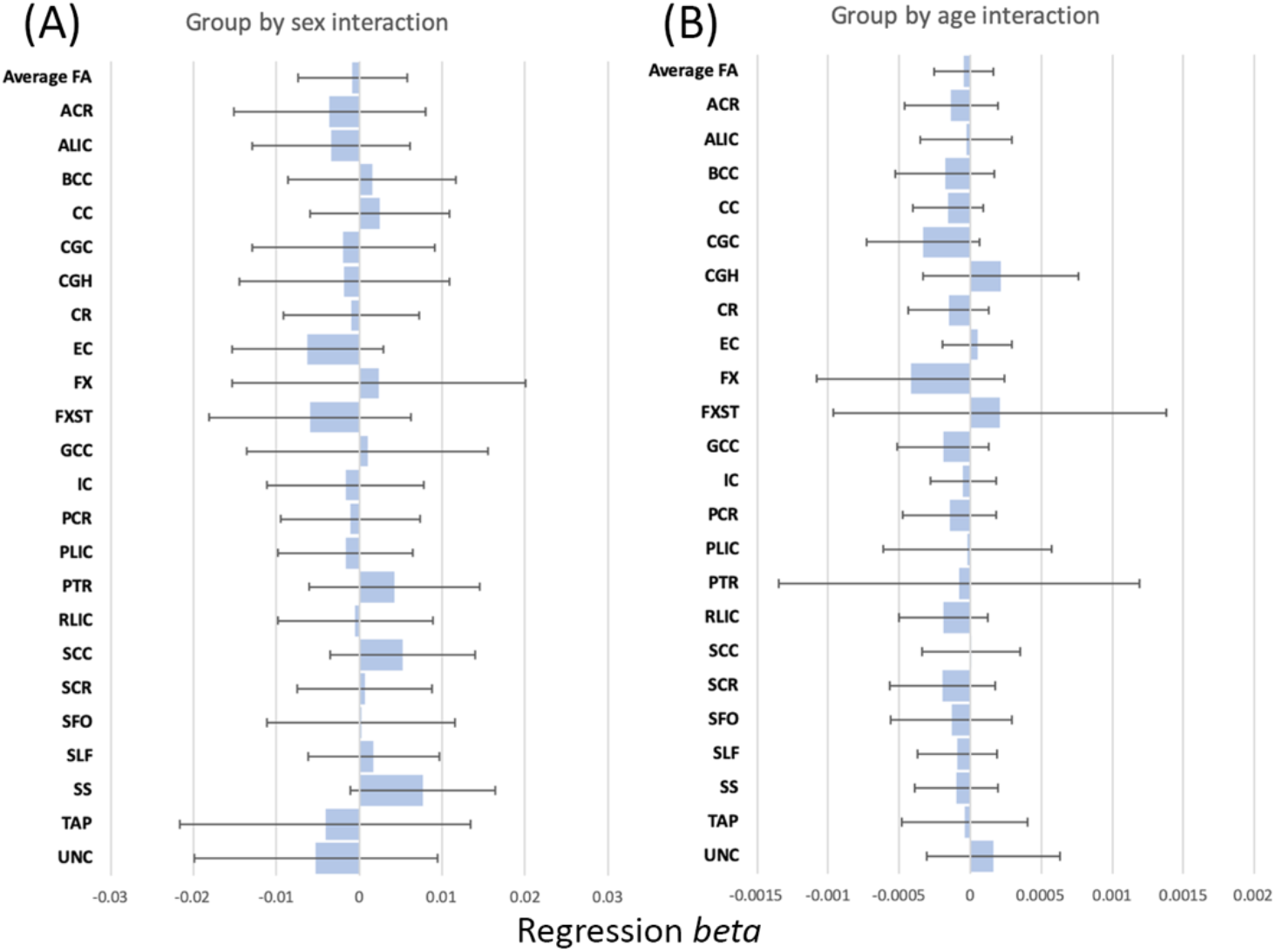
Interactions: (a) group-by-sex interaction effects and (b) group-by-age interaction effects. Shown are regression bs for 23 ROIs and average FA. Dark orange bars indicate significance (*p*<0.0021) and light orange bars indicate marginally significant results (0.05>*p*>0.0021). Error bars are 95% CI.

**Supplementary Figure 5.**
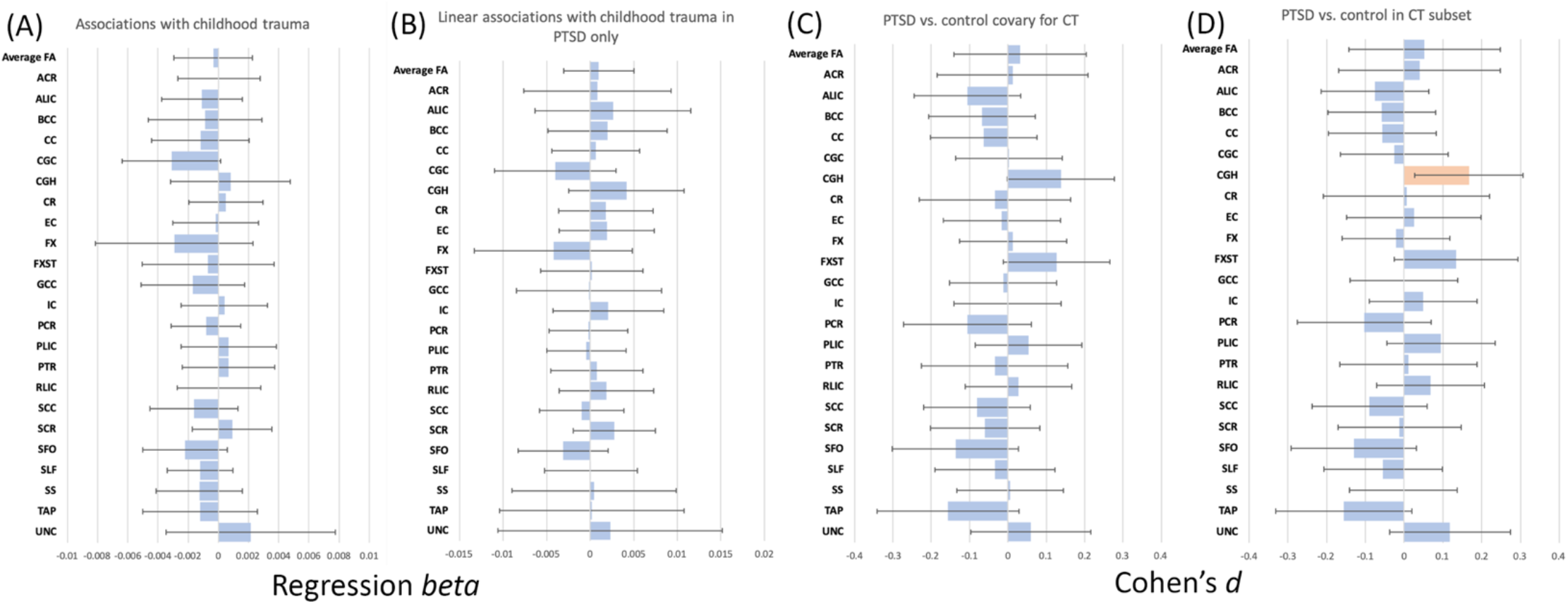
Results of childhood trauma (CT) analyses: (a) linear regression with CT (coded 0=none, 1=1 type, 2=2 or more types of trauma) with sex, age, and age^2^ in the model; (b) linear regression with CT in PTSD group only; (c) PTSD vs. control group differences when covarying for CT; (d) PTSD vs. control differences in the subset of participants with CT, WITHOUT CT in model. Shown are Cohen’s *d* or unstandardized regression bs for 23 ROIs and average FA. Dark orange bars indicate significance (*p*<0.0021) and light orange bars indicate marginally significant results (0.05>*p*>0.0021). Error bars are 95% CI.

**Supplementary Figure 6.**
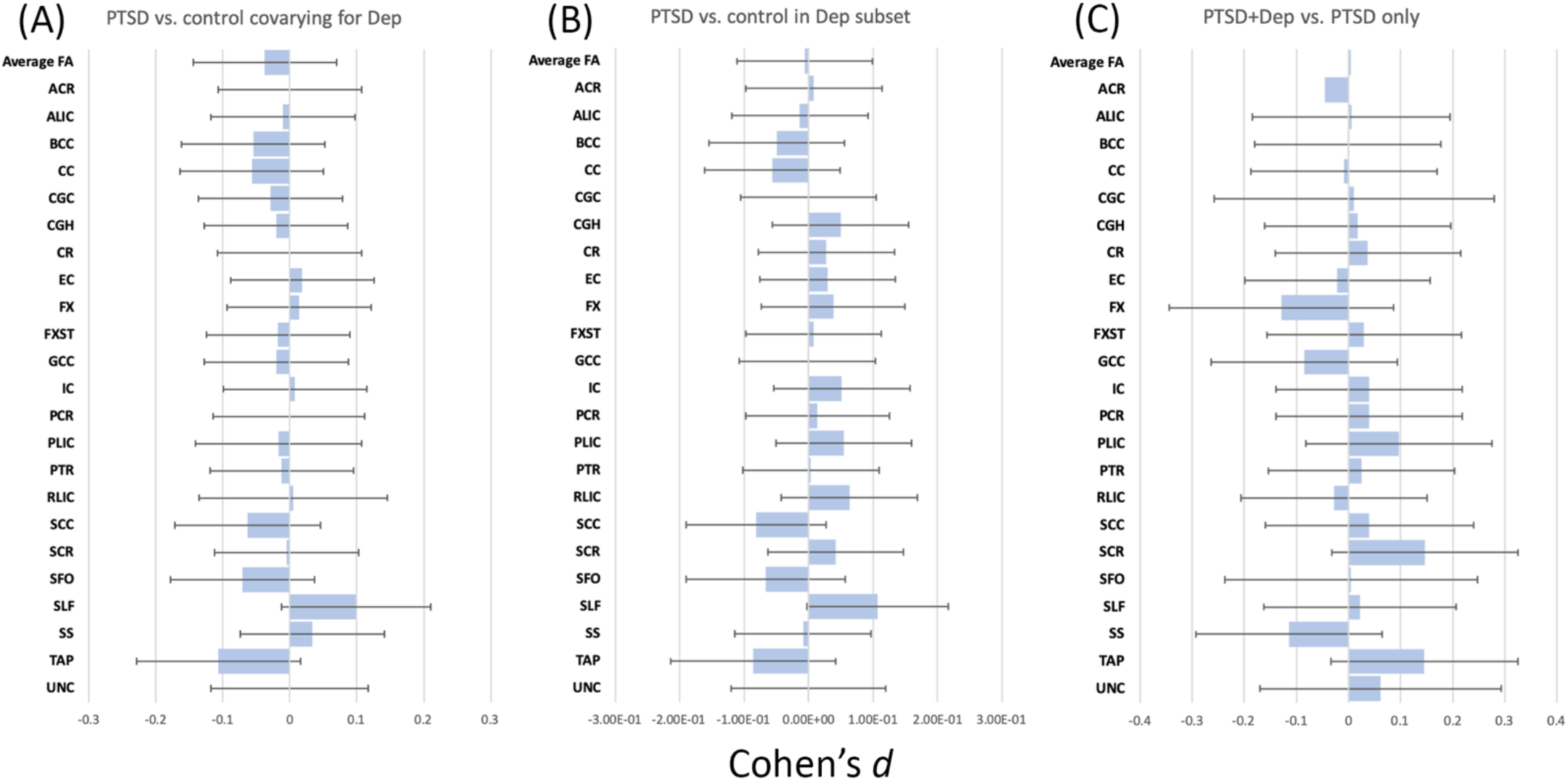
Results of depression analyses: (a) PTSD vs. control group differences when covarying for depression; (b) PTSD vs. control differences in the subset of participants with depression, WITHOUT depression in model; (c) PTSD+depression vs. PTSD only differences. Shown are Cohen’s *d* for 23 ROIs and average FA. Dark orange bars indicate significance (*p*<0.0021) and light orange bars indicate marginally significant results (0.05>*p*>0.0021). Error bars are 95% CI.

**Supplementary Figure 7.**
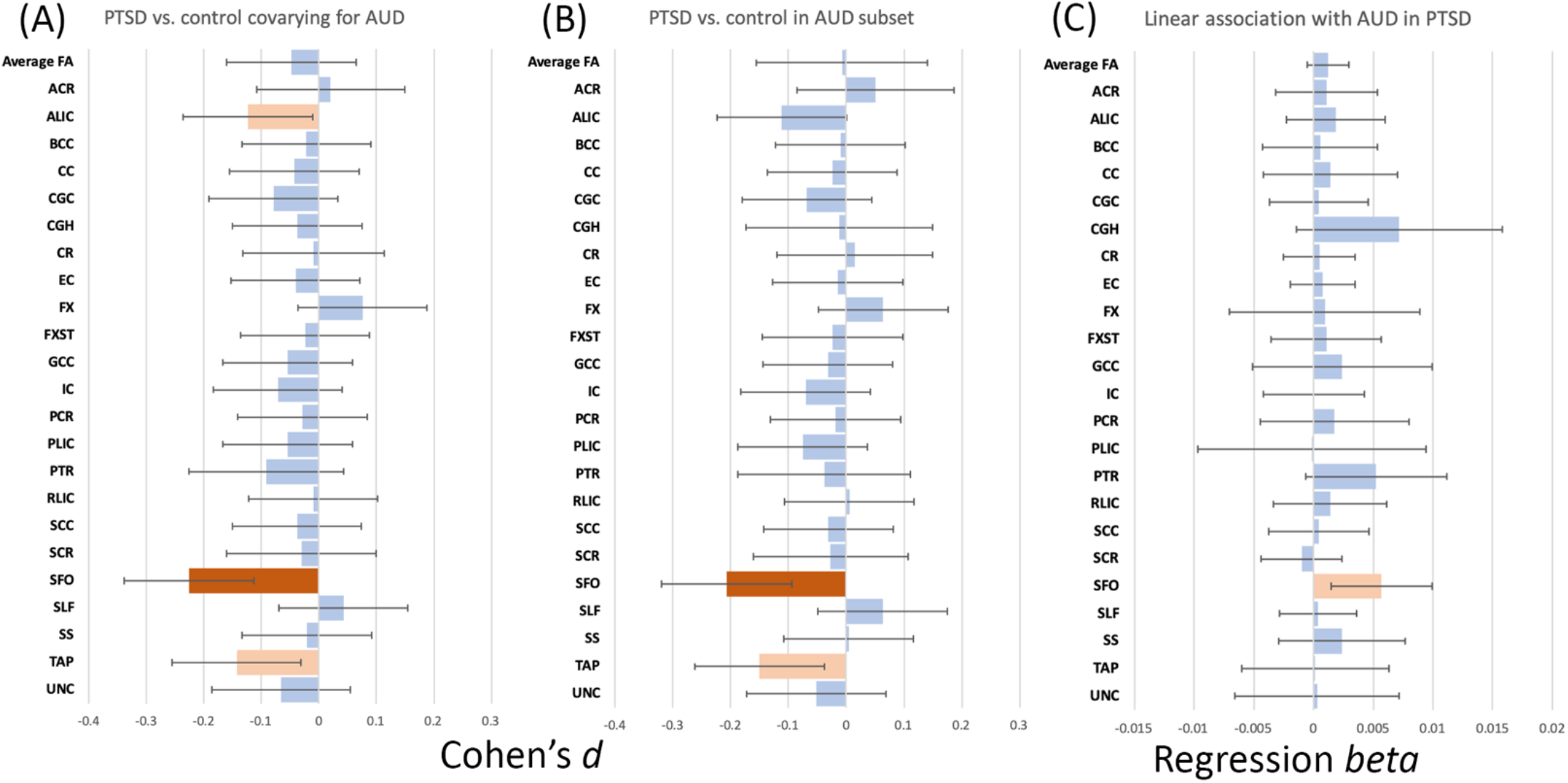
Results of alcohol use disorder (AUD) analyses: (a) PTSD vs. control group differences when covarying for AUD; (b) PTSD vs. control differences in the subset of participants with AUD, WITHOUT AUD in model; (c) linear association with AUD in PTSD group. Shown are Cohen’s *d* or unstandardized regression bs for 23 ROIs and average FA. Dark orange bars indicate significance (*p*<0.0021) and light orange bars indicates marginally significant results (0.05>*p*>0.0021). Error bars are 95% CI.

**Supplementary Figure 8.**
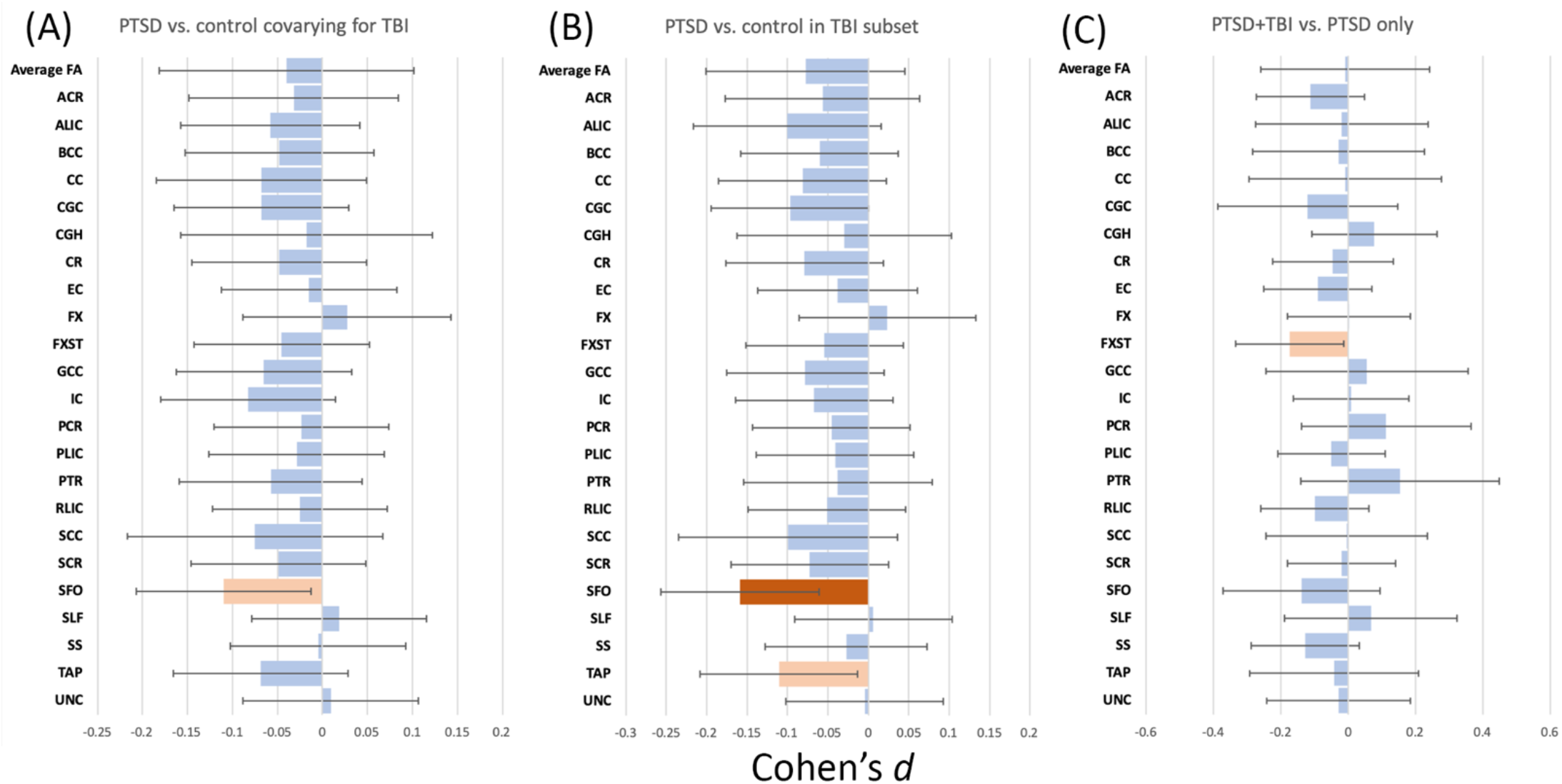
Results of traumatic brain injury (TBI) analyses: (a) PTSD vs. control group differences when covarying for TBI; (b) PTSD vs. control differences in the subset of participants with TBI, WITHOUT TBI in model; (c) PTSD+TBI vs. PTSD only differences. Shown are Cohen’s *d* for 23 ROIs and average FA. Dark orange bars indicate significance (*p*<0.0021) and light orange bars indicates marginally significant results (0.05>*p*>0.0021). Error bars are 95% CI.

**Supplementary Figure 9.**
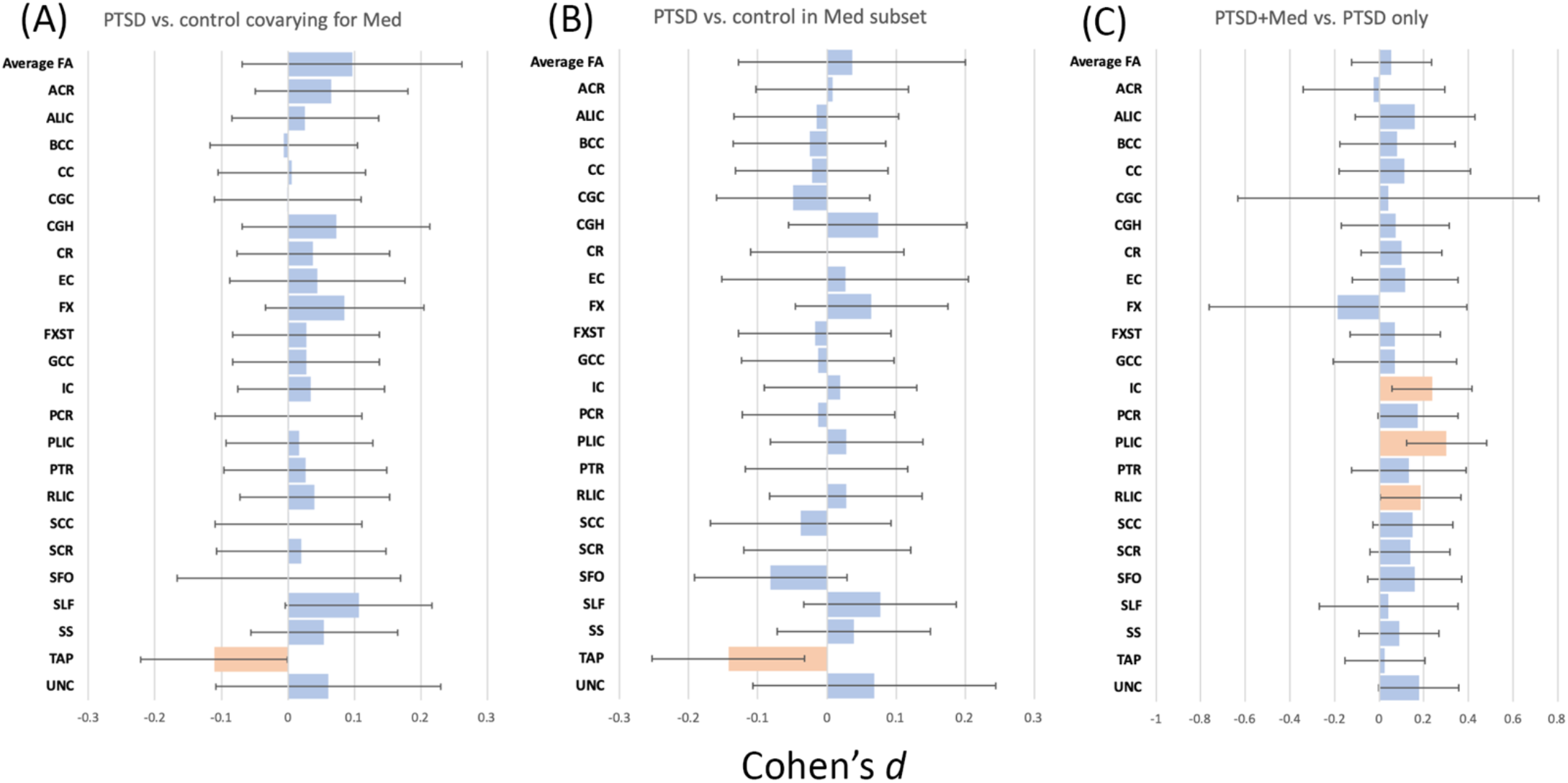
Results of medication analyses: (a) PTSD vs. control group differences when covarying for psychotropic medication use; (b) PTSD vs. control differences in the subset of participants with medication, WITHOUT medication in model; (c) Medicated PTSD vs. unmedicated PTSD differences. Shown are Cohen’s *d* for 23 ROIs and average FA. Dark orange bars indicate significance (*p*<0.0021) and light orange bars indicate marginally significant results (0.05>*p*>0.0021). Error bars are 95% CI.

**Supplementary Figure 10.**
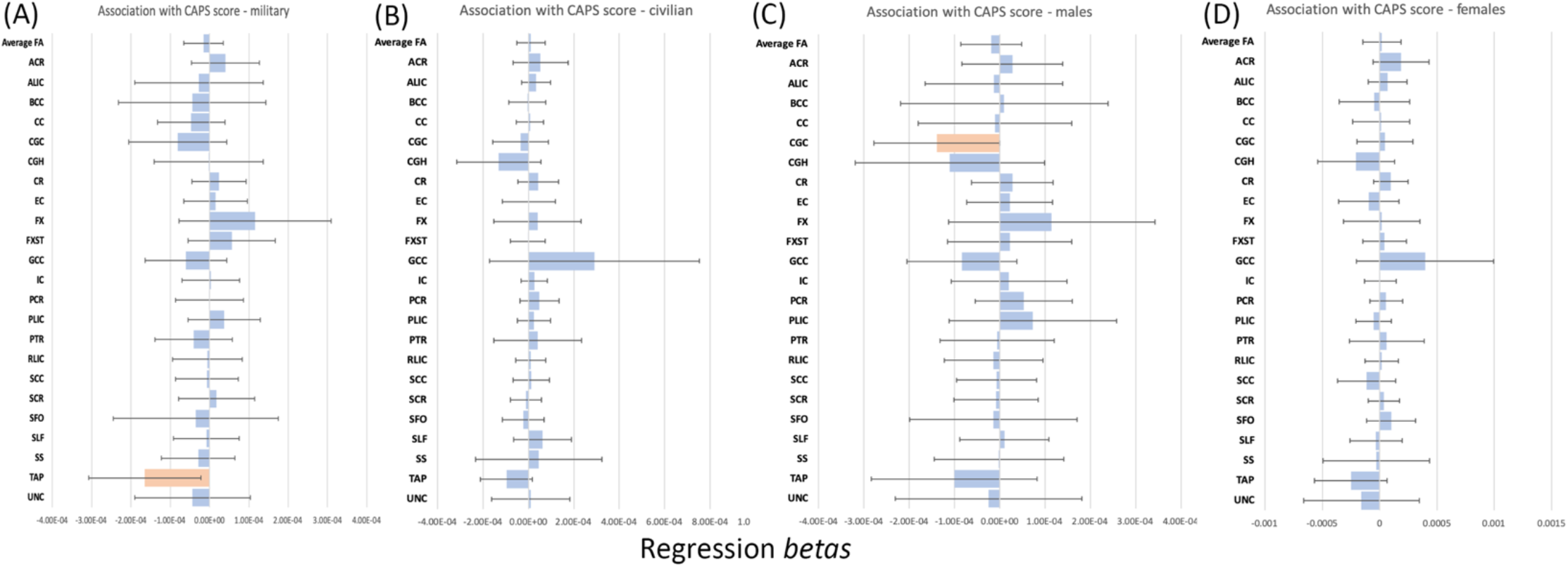
Linear association with CAPS within subgroups: (a) military, (b) civilian, (c) males, (d) females. Shown are unstandardized regression bs for 23 ROIs and average FA. Dark orange bars indicate significance (*p*<0.0021) and light orange bars indicate marginally significant results (0.05>*p*>0.0021). Error bars are 95% CI.

**Supplementary Figure 11.**
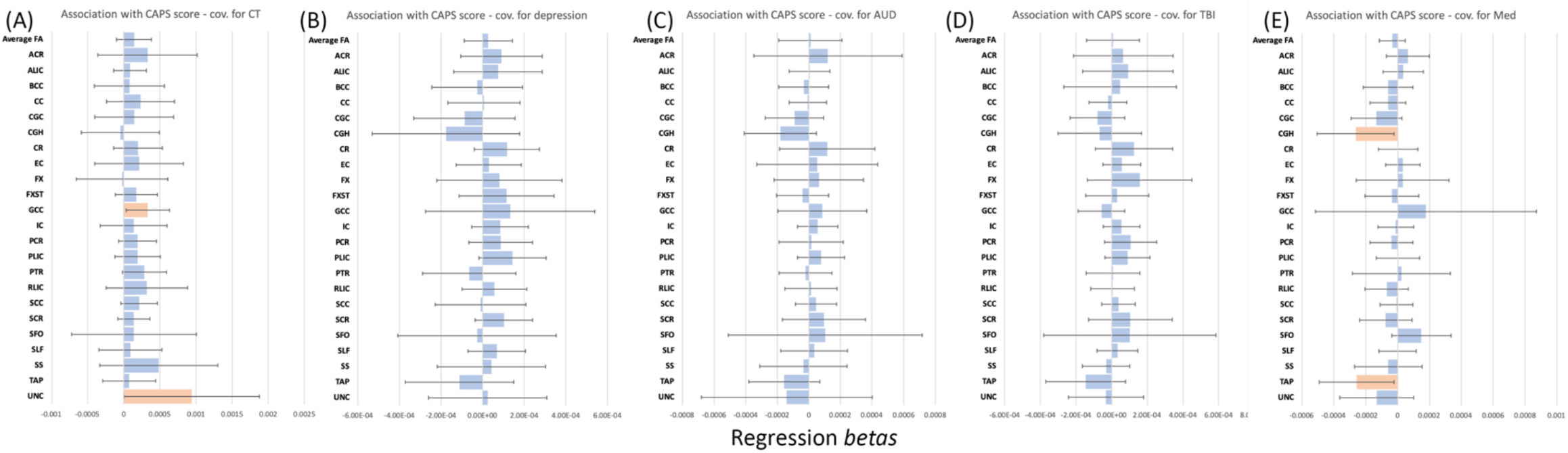
Linear association with CAPS including potentially confounding variables: (a) childhood trauma (0=none, 1=1 type, 2=2 or more types), (b) depression (0=no, 1=yes), (c) alcohol use disorder (0=none, 1=alcohol abuse, 2=alcohol dependence), (d) TBI (0=no, 1=yes), and (e) psychotropic medication use (0=no, 1=yes). Shown are unstandardized regression bs for 23 ROIs and average FA. Dark orange bars indicate significance (*p*<0.0021) and light orange bars indicate marginally significant results (0.05>*p*>0.0021). Error bars are 95% CI.

**Supplementary Figure 12.**
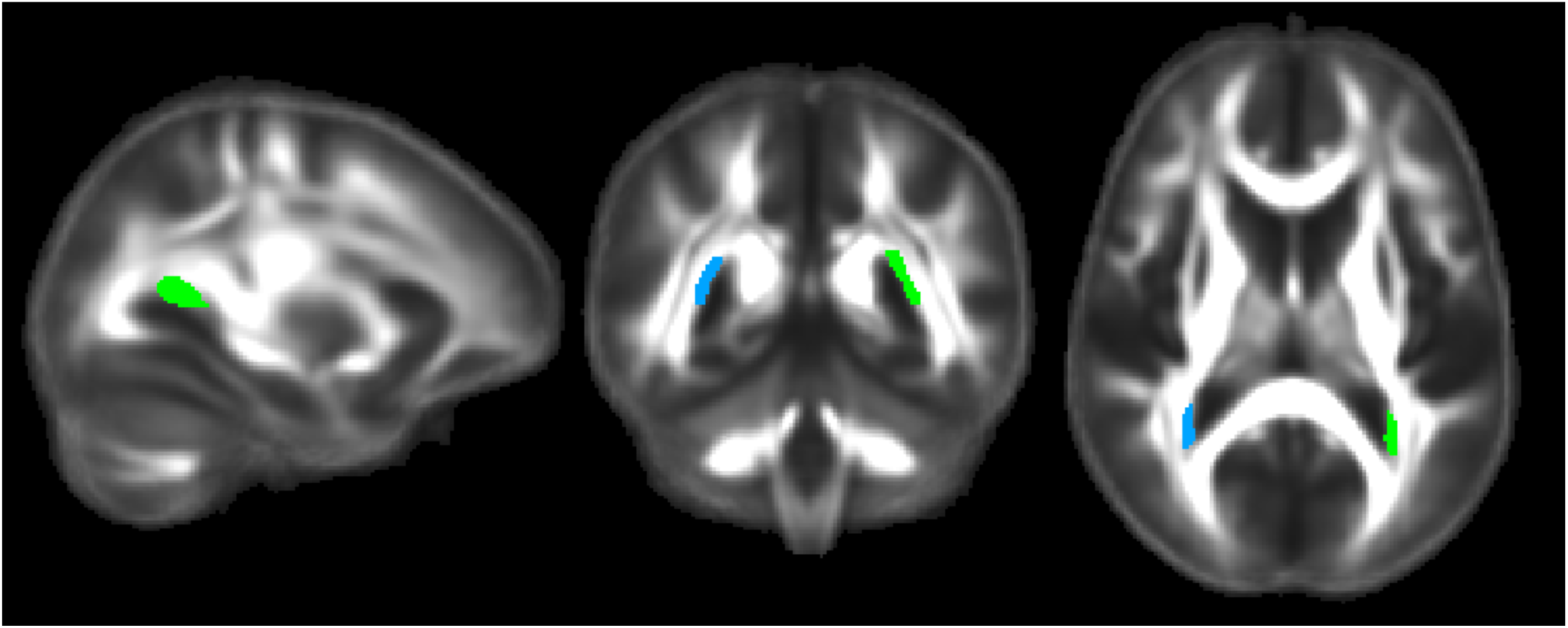
Tapetum displayed on the ENIGMA template FA. The left tapetum (green) and right tapetum (blue) ROIs are displayed. Left in image is right in brain.

### Supplementary Note 1. Further detail of ENIGMA-DTI protocols

Details on scanner and acquisition parameters are provided in **Supplementary Table 2**. Preprocessing, including eddy current correction, EPI induced distortion correction, and tensor fitting, was carried out at each site. Image analysis was conducted at each site using tract-based spatial statistics (TBSS) as part of FSL software^1^. Individual subject FA maps were aligned to the custom ENIGMA-DTI FA template derived from 400 adult participants scanned across four sites designed for optimal multi-site harmonization^2^. FA voxels were then projected onto the ENIGMA-DTI template skeleton. This creates a unique FA skeleton in the same space for each individual in each cohort. To minimize effects of residual registration misalignment, the regions of interests were consistent in size across sites and the skeletonization procedure was performed individually at each site to minimize any site-specific residual misalignment. The same projection used for the FA images also projects the non-FA (mean, axial, and radial) images onto the skeleton. Voxels along the individual skeletons were averaged across white matter ROIs. A total of 25 bilateral ROIs were delineated based on the JHU WM atlas, an established WM parcellation derived using deterministic tractography^3^. A whole-brain WM skeleton was defined according to the tract-based spatial statistic methodology^1^ and ROI-averaged measures of FA, MD, AD and RD were then calculated by averaging each of these voxel measures over all skeleton voxels encapsulated by a particular ROI. This ensured that voxels at the periphery of a fiber bundle, where residual registration misalignment is typically maximal, were excluded from the ROI average. In other words, ROI averaging was performed based on the core of each fiber bundle, as defined by the WM skeleton. The multi-subject JHU white matter parcellation atlas^3^ was used to parcellate regions of interest from the ENIGMA template in MNI space, with updated label identification to correct an earlier atlas error^4^. A total of eighteen bilateral white matter ROIs were extracted from the skeletonized FA images and averaged (the corticospinal tract was ignored as prior reports have shown it to have poor reliability). The table below lists 24 ROIs (some partially overlapping) that were extracted from the skeletonized images, including 5 midsagittal regions (no lateralized components), and 18 lateralized regions (left and right are averaged to obtain bilateral FA). The overall average FA values were calculated by averaging values for the entire white matter skeleton. ENIGMA-DTI QA/QC protocol consists of visual inspection of the images before and after registration to the ENIGMA template, as well as calculating the average skeleton projection distance. The distance of voxel projection to the ENIGMA skeleton can assess the registration quality between individual images and ENIGMA-DTI template. Higher projection distance may indicate problems with aligning individual brain to the template. After ROI extraction, histograms of FA and diffusivity measures are computed for each ROI.

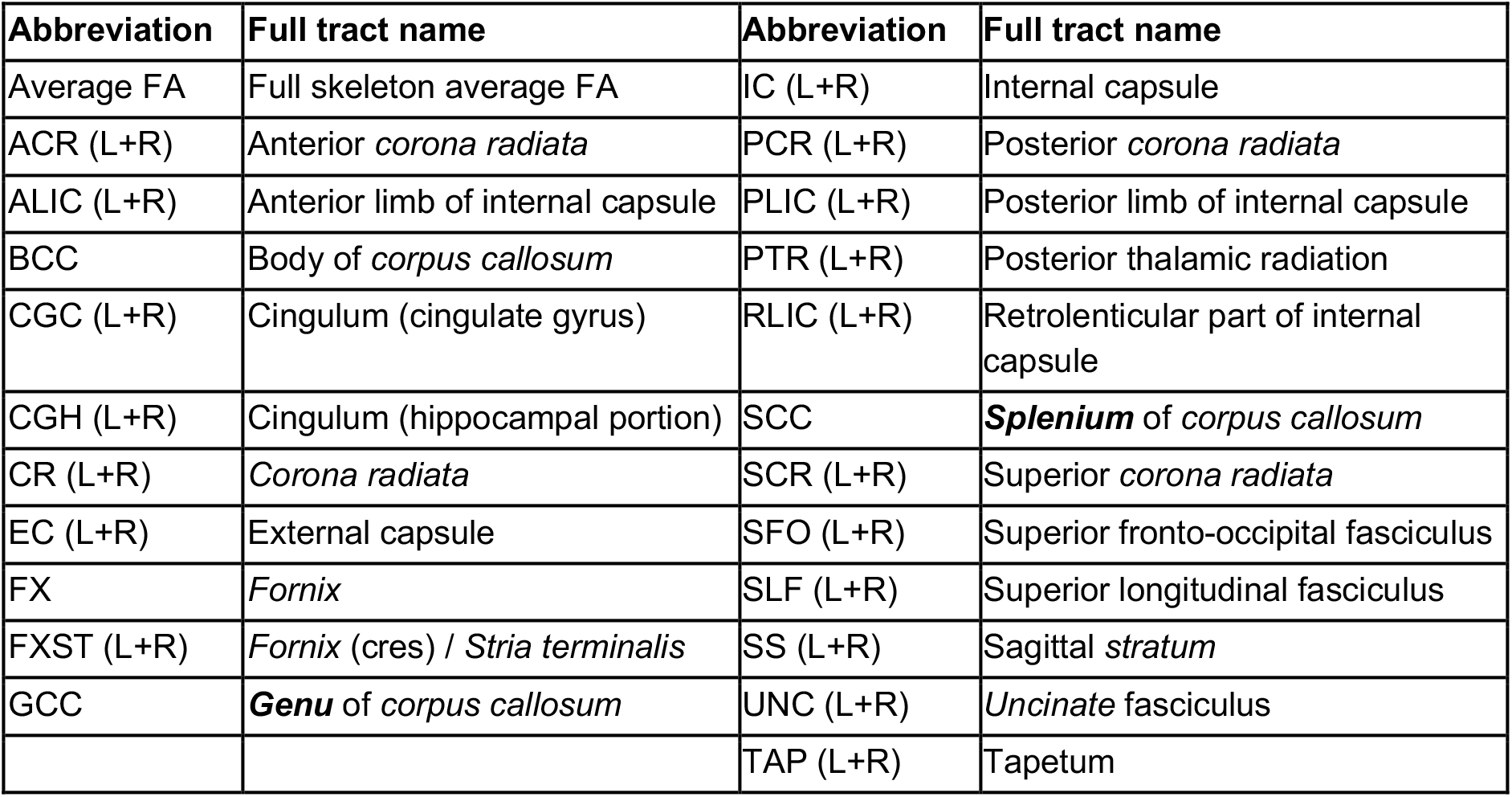

| Abbreviation | Full tract name | Abbreviation | Full tract name |
| --- | --- | --- | --- |
| Average FA | Full skeleton average FA | IC (L+R) | Internal capsule |
| ACR (L+R) | Anterior *corona radiata* | PCR (L+R) | Posterior *corona radiata* |
| ALIC (L+R) | Anterior limb of internal capsule | PLIC (L+R) | Posterior limb of internal capsule |
| BCC | Body of *corpus callosum* | PTR (L+R) | Posterior thalamic radiation |
| CGC (L+R) | Cingulum (cingulate gyrus) | RLIC (L+R) | Retrolenticular part of internal capsule |
| CGH (L+R) | Cingulum (hippocampal portion) | SCC | ***Splenium*** of *corpus callosum* |
| CR (L+R) | *Corona radiata* | SCR (L+R) | Superior *corona radiata* |
| EC (L+R) | External capsule | SFO (L+R) | Superior fronto-occipital fasciculus |
| FX | *Fornix* | SLF (L+R) | Superior longitudinal fasciculus |
| FXST (L+R) | *Fornix* (cres) / *Stria terminalis* | SS (L+R) | Sagittal *stratum* |
| GCC | ***Genu*** of *corpus callosum* | UNC (L+R) | *Uncinate* fasciculus |
|  |  | TAP (L+R) | Tapetum |

### Supplementary Note 2. Details on depression inventories

As depression was assessed using a range of scales, we used published clinical cutoffs to establish a categorical depression variable. For the Beck Depression Inventory (BDI), the cutoff for depression was >13^5^. For the Geriatric Depression Scale (GDS), the cutoff for depression was >4^6^. For the Center for Epidemiologic Studies – Depression scale (CES-D), the cutoff for depression was >15^7^. For the Hamilton Depression Inventory (HAM-D), the cutoff for depression was >8^8^. For the Hospital Anxiety and Depression Scale – Depression (HADS-D), the cutoff for depression was >7^9^. The Structured Clinical Interview for DSM-V (SCID) yields a categorical variable not a score so it did not need to be converted. The cite contributing DASS scores did not separate the depression from anxiety variables, so it was not included in the depression analyses.

### Supplementary Note 3. Subgroup analyses

Comparing PTSD to control within the female participants (259 PTSD vs. 316 controls), we did not find any significant effects. Within the male participants (1022 PTSD vs. 1120 controls) we found marginally lower FA in the bilateral tapetum and SFO (*d*=−0.13, *p*=0.0052; *d*=−0.14, *p*=0.011), and lower FA in PTSD in the left tapetum (*d*=−0.15, *p*=0.0007), along with marginally higher RD in the left tapetum (*d*=0.010, *p*=0.037). Within the military cohorts (988 PTSD vs. 1085 controls), we found marginally lower FA and AD in the bilateral tapetum and right SFO (FA: *d*=−0.11, *p*=0.015; *d*=−0.12, *p*=0.011; bilateral tapetum AD: *d*=−0.10, *p*=0.049) and lower FA in PTSD in the left tapetum (*d*=−0.16, *p*=0.00066). Among civilian cohorts (409 PTSD vs. 518 controls), we did not find any significant or marginally significant results. Examining the site effects in **Figure 2**, several civilian cohorts did have large effect sizes for the tapetum, so the lack of results may be due in part to the significantly smaller civilian sample size compared to military. These results can be seen in **Supplementary Figure 3.** Lastly, as results seemed to be strongest in the males-only and military-only analyses, we further examined military females and military males separately. Among military females (44 PTSD vs. 90 controls), we did not find any significant results. Among military males (693 PTSD vs. 708 controls), we found marginally lower FA in the left tapetum (*d*=−0.14, *p*=0.012).

We similarly examined linear association with CAPS-4 score in these subgroups. In female PTSD participants there were no significant effects. In male PTSD participants, we marginally lower FA with higher CAPS-4 in the cingulum (b=−0.00014, *p*=0.050). In civilian PTSD participants we did not find any significant effects. In military PTSD participants, we found marginally lower FA with higher CAPS-4 in the bilateral and right tapetum (b=−0.00016, *p*=0.024; b=−0.00020, *p*=0.026, respectively). These results can be seen in **Supplementary Figure 10.**

### Supplementary Note 4. Interaction analyses

Examining a group-by-age interaction variable with age, age^2^, sex, and group as covariates, we did not find any significant or marginally significant results. We similarly did not detect any interactions between group and sex. These results can be seen in **Supplementary Figure 4.**

### Supplementary Note 5. Full results covarying for depression, alcohol use disorders, traumatic brain injury, or psychotropic medication use

#### Potentially confounding variables

For all of the following analyses, we did not include sites with fewer than 10 subjects per cell.

We did not find any significant associations with CT covarying for age, age^2^, and sex. Including CT in a model comparing PTSD and control (367 PTSD vs. 598 controls) also did not yield significant results. Comparing PTSD and controls in the same reduced sample, without CT in the model showed higher FA in the CGH (*d*=0.17, *p*=0.019), suggesting that the decrease in power was driving differences between these and the main analyses. Lastly, we examined linear associations with CT in the PTSD group only, finding no significant associations. For this analysis, there were 57 PTSD with no CT, 48 PTSD with 1 type of CT exposure, and 179 PTSD with 2 types of CT exposure across 6 sites.

Including a binary depression variable as a covariate in PTSD vs. control comparisons (696 PTSD vs. 825 controls), we found marginally lower FA in the left tapetum (*d*=−0.15, *p*=0.0090). In this reduced sample (N=1521) without depression in the model, results were similar (left tapetum *d*=−0.12, *p*=0.048). Comparing PTSD+Dep to PTSD only (304 PTSD+Dep vs. 225 Dep only, from 12 sites), we did not find any significant effects.

Including AUD (coded as 0=no alcohol use disorder, 1=abuse, 2=dependence) as a covariate in PTSD vs. control comparisons (691 PTSD vs. 623 controls), we found significantly lower FA in the right SFO (*d*= −0.23, *p*=0.000083), and marginally lower FA and higher RD in the tapetum (bilateral *d*=−0.14, *p*=0.012; left *d*= −0.16, *p*=0.0061; left tapetum RD *d*=0.12, *p*=0.037) and ALIC (bilateral *d*=−0.12, *p*=0.031; left *d*=−0.15, *p*=0.020). In this reduced sample (N=1314) without AUD in the model, results were essentially the same. Examining linear associations with AUD in the PTSD group only (419 PTSD with no AUD, 77 PTSD + alcohol abuse, 113 PTSD + alcohol dependence, across 10 sites), we found marginally higher FA in PTSD+AUD in the right SFO (b=0.0057, *p*=0.0082).

Including a binary TBI variable in PTSD vs. control comparisons (849 PTSD vs. 1016 controls), we found marginally lower FA in the left tapetum (*d*=−0.12, *p*=0.015) and right SFO (*d*=−0.11, *p*=0.027). The analysis in this reduced sample (N=1865) without TBI in the model yielded similar results with the addition of marginally lower FA in the bilateral tapetum (*d*=−0.11, *p*=0.026). Comparing PTSD+TBI to PTSD only (462 PTSD+TBI vs. 270 PTSD only, across 9 sites), we found marginally lower FA in the PTSD+TBI group in the fornix/stria terminalis (*d*=−0.17, *p*=0.033).

Including a binary variable for psychotropic medication use in PTSD vs. control comparisons (713 PTSD vs. 679 controls), we found marginally lower FA in the bilateral and left tapetum (bilateral *d*=−0.11, *p*=0.050; left *d*=−0.014, *p*=0.013). The analysis in this reduced sample (N=1392) without medication in the model yielded similar results. Comparing medicated PTSD to unmedicated PTSD (268 med-PTSD vs. 221 unmed-PTSD, across 5 sites), we found marginally higher FA in the medicated PTSD group in the PLIC, IC, and RLIC (*d*=0.30, *p*=0.0010; *d*=0.24, *p*=0.0097; *d*=0.19, *p*=0.042, respectively).

These results can be seen in **Supplementary Figures 5–9.** The similarities between the group comparisons with the added covariate (depression, AUD, or TBI) and the group comparisons without that covariate in the subset of the sample with that information indicate that the decrease in sample size is affecting our main tapetum result more than the addition of the covariate itself.

#### Confounding variables with PTSD severity

We similarly ran analyses examining linear associations with CAPS-4 score including potentially confounding variables. Covarying for CT, we found marginally higher FA with higher CAPS-4 score in the uncinate and genu (b=0.00094, *p*=0.050; b=0.00033, *p*=0.030). Covarying for depression, we did not find any significant associations with CAPS-4 score. Covarying for AUD, we did not find any significant associations with CAPS-4 score. Covarying for TBI, we did not find any significant association with CAPS-4 score. Covarying for psychotropic medication use, we found marginally lower FA with higher CAPS-4 score in the tapetum and hippocampal cingulum (b=−0.00026, *p*=0.032; b=−0.00026, *p*=0.032). These results can be seen in **Supplementary Figure 11.**

